# A meningeal circadian clock and brain state co-ordinately enhance macromolecular clearance during wake

**DOI:** 10.64898/2025.12.25.696495

**Authors:** Talya Goble, Tanushree Kundu, Léticia S. Meyer, Dimitra Klinaki, Thomas A. Hawkins, Jason Rihel

## Abstract

The removal of brain waste is thought to be regulated by the circadian clock and sleep-wake cycle. A meningeal scavenger cell population, called mural Lymphatic Endothelial Cells (muLECs), uptake macromolecules from the cerebrospinal fluid (CSF) and are poised to clear soluble brain waste. Whether CSF clearance is regulated across the 24-hour day is unknown. Using a quantitative, *in vivo* assay in zebrafish, we show that the capacity of muLECs to remove macromolecules is highest during the waking day. The muLEC protein uptake is regulated by a cell-autonomous circadian oscillator that drives rhythmic protein levels of Mannose Receptor 1a (Mrc1a); in contrast, lipoprotein uptake increases as a function of time spent awake. Thus, circadian and brain state cues independently maximize muLECs’ function during wake. Consistently, muLECs are required for normal recovery after the heightened metabolic demand of a seizure, suggesting that muLEC clearance contributes to both normal and disrupted brain homeostasis.

**Impact Statement:** A meningeal scavenger cell population called Mural Lymphatic Endothelial Cells preferentially clear macromolecules from the CSF during the waking day, and this uptake is regulated independently by brain state and circadian cues in a substrate-specific manner.

## Introduction

The mechanisms that maintain fluid homeostasis and clearance of waste macromolecules from the brain remain incompletely understood. Current frameworks propose that brain waste clearance involves the movement and subsequent excretion of soluble waste via fluid exchange between the interstitial fluid (ISF) of the brain and the cerebrospinal fluid (CSF) of the encasing leptomeninges along with intracellular degradation of waste by resident parenchymal and meningeal cell populations (Neumann et al, 2008; Liu et al, 2022; Zhan et al, 2025). In 2017, a new scavenger cell type was discovered that resides in the meninges of zebrafish termed Mural Lymphatic Endothelial Cells (muLECs; also termed brain LECs and Fluorescent Granular Perithelial cells) (Bower et al, 2017; Galanternik et al, 2017; van Lessen et al, 2017), and subsequently a similar cell type has been described in mammals (Shibata-Germanos et al, 2020; Siret et al, 2022). Similar to other LECs, muLECs arise from vascular endothelium and rely on lymphatic developmental signals (Bower et al, 2017; Galanternik et al, 2017; van Lessen et al, 2017). Uniquely, however, muLECs do not form lumenised vessels, instead forming a loosely connected meshwork of cells that closely associate with meningeal blood vessels covering the adult zebrafish brain (van Lessen et al, 2017). Although the physiological functions of muLECs are yet unknown, these cells are poised as CSF scavengers to clear metabolic waste from the brain. For example, muLECs readily endocytose a variety of macromolecules spanning proteins, lipoproteins, liposomes, and polysaccharides, with a higher uptake efficiency than brain-resident microglia (Huisman et al, 2021). Recent work has demonstrated that muLEC development is sensitive to, and regulated by, neuronal activity (Li et al, 2025), hinting at a functional association between the brain and this meningeal cell population.

Over the past decade, numerous studies have reported that brain clearance pathways are regulated by both behavioural state and circadian cues, although there is some discrepancy as to whether this drives peak clearance preferentially during the sleep or wake period. CSF oscillations in the ventricles that are dependent on sleep stages have been observed in humans (Fultz et al, 2019) and birds (Ungurean et al, 2023) and are thought to drive fluid influx into the brain. In mice, CSF influx has been reported to be higher during sleep (Xie et al, 2013), potentially driven by sleep-associated changes in arterial pulsations (Bojarskaite et al, 2023) and circadian regulation of Aquaporin-4 water channels during the subjective rest period (Hablitz et al, 2020). Clearance of exogenous tracers from the brain has been shown to be elevated during sleep (Xie et al, 2013; Eide et al, 2021), whereas lymphatic drainage of CSF from the head cavity via meningeal lymphatic vessels was higher during the subjective active period (Hablitz et al, 2020). Microglia, the brain-resident phagocytes, show sleep- and circadian-regulation in morphology and uptake capacity of amyloid β (Liu et al, 2019; Clark et al, 2022; Parhizkar et al, 2023). However, recent work has reported preferential tracer clearance during wakefulness (Miao et al, 2024), challenging the notion that brain clearance is enhanced during sleep and may in fact be highest during wake. Nevertheless, these data collectively demonstrate brain clearance is dynamic across the day-night cycle. Given that muLECs are poised to remove waste from the brain, we hypothesized that their function may likewise show dynamic regulation over the 24-hour day-night cycle.

Here, we developed an *in vivo*, quantitative method to measure muLEC CSF solute clearance over the day-night cycle in zebrafish larvae and show that clearance of multiple substrate classes is highest during the waking day. Experimental disentanglement of light, circadian, and sleep-wake cues surprisingly revealed that while muLEC-uptake capacity for both protein and low-density lipoprotein (LDL) peaks during the day-time, these are independently regulated by a muLEC-autonomous circadian oscillator and sleep-wake state, respectively. Perturbing muLEC function slowed behavioural recovery following a seizure, suggesting that muLECs contribute to maintaining solute homeostasis during periods of high metabolic demand.

## Results

### muLEC morphology is dynamic over the day-night cycle

If muLEC function changes across the 24-hour day-night cycle, we reasoned that this regulation may be detectable as alterations in cellular morphology during the day versus night. To test this, we first raised *Tg(flt4:mCitrine)* embryos, which have fluorescently labelled muLECs (van Impel et al, 2014) and other lymphatic vasculature, to 6 days post fertilisation (6dpf) on opposing 14hr:10hr light-dark cycles to yield developmentally matched larvae in different phases of their day-night cycle. At 6 dpf, muLECs form bilateral loops in the developing meninges dorsal to the optic tectum (Figure 1a) and harbour distinctive, round intracellular inclusions of varying sizes and filopodial-like structures (Figure 1d; S1a; S1g). During the day, the total volume (Figure 1b, p=0.008) and surface area (Figure S1c, p=0.0007) of muLECs were significantly bigger compared to at night. Curiously, these daytime size changes were also accompanied by a reorganization of muLEC interactivity, with significantly more disconnections between muLECs during the day versus the night (Figure 1c, p=0.049), although neither the width (Figure S1e, p=0.15) nor length (Figure S1f, p=0.27) of remaining connections were altered (Figure S1b). In addition, the muLECs had a trend towards more filopodia (Figure 1e) and significantly more intracellular inclusions (Figure 1f, p=0.02) during the day than at night, although there were no changes in either filopodial length (Figure S1d, p=0.45) or average inclusion size (Figure 1g, p=0.24). As an increased number of intracellular inclusions likely represent increased macromolecular internalization, these results suggest that muLEC scavenger activity is higher during the waking day than at night.

**Figure 1:**
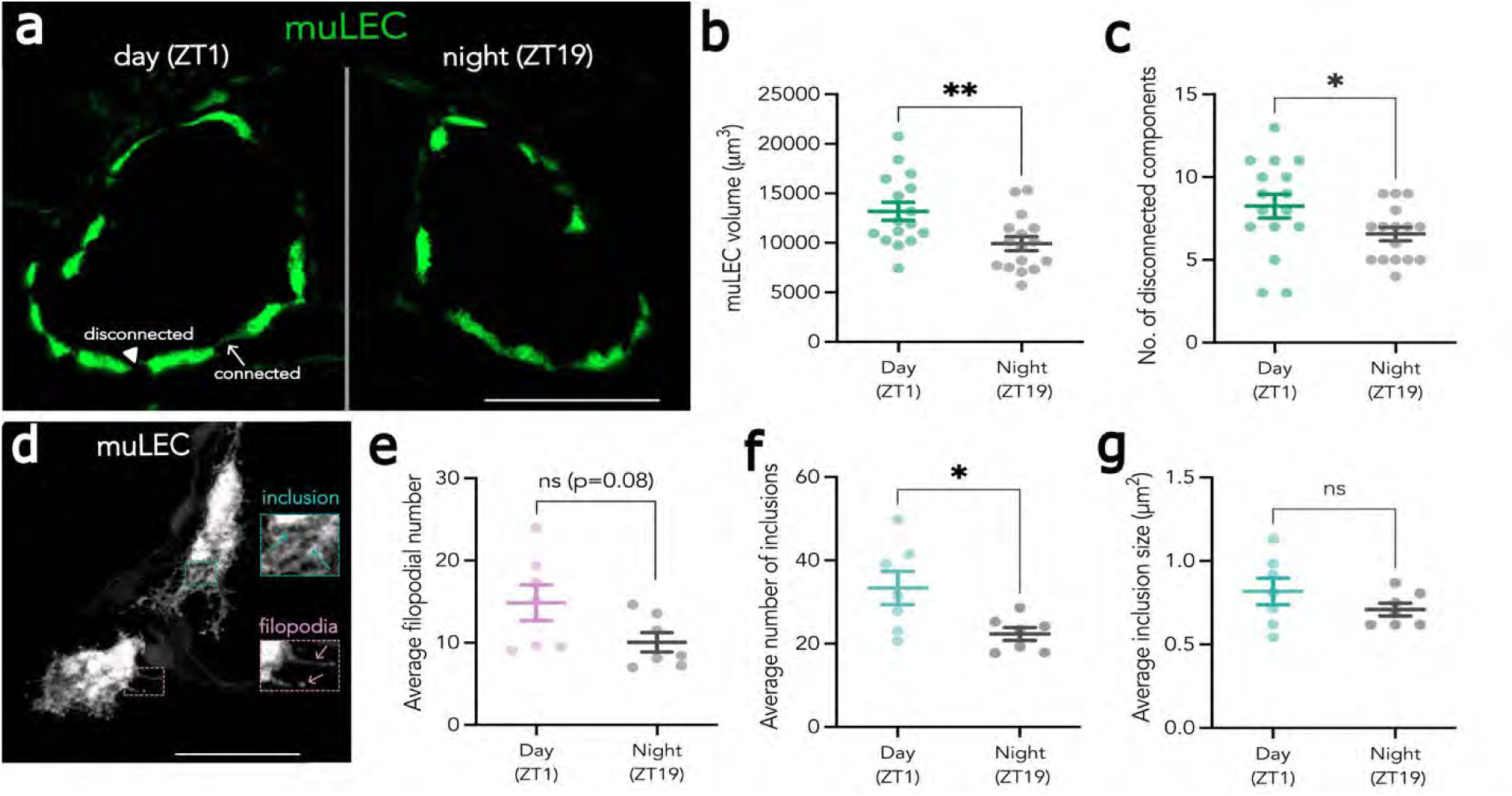
muLEC morphology is dynamic over the day-night cycle. **a**) Representative maximum projection of a muLEC tectal loop (green) in 6dpf Tg*(flt4:mCitrine)* larvae during the day (ZT1, left) versus night (ZT19, right). Connected (arrow) and disconnected (arrowhead) muLECs are highlighted. **b**) Quantification of total muLEC volume within a tectal loop during the day (n=16) and night (n=16). **c**) Quantification of muLEC disconnectedness in the day (n=16) and night (n=16). **d**) High-resolution muLEC imaging reveals fluorescent negative inclusions (blue arrows) and filopodia-like protrusions (pink arrows). **e**) Average number of muLEC filopodial protrusions per cell per larva in the day (n=7) and night (n=7). (**f** & **g**) Average size (f) and number (g) of intracellular muLEC inclusions, day (n=7) and night (n=7). Data are mean ± SEM. Each dot represents one larva. Statistical differences were determined by one-way ANOVA (two-tailed, b-g). ^ns^ p>0.05; * p< 0.05; ** p<0.01. Abbreviations: ZT, Zeitgeber Time. Scale bar = 100µm (a), 20µm (b).

### muLEC-uptake of diverse macromolecules is higher during the day

To investigate whether diurnal changes in muLEC morphology relate to a difference in uptake capacity, we developed a quantitative assay to measure muLEC-uptake rates *in vivo*. Dorsally mounted *Tg(flt4:mCitrine)* larvae were co-injected into the hindbrain ventricle with the pH-sensitive dye, pHrodo-Red avidin (hereafter termed pHrodo-avidin), which is readily internalized by muLECs into acidified compartments, and a 2MDa Alexa-fluor, 488 dextran (hereafter termed 2MDa-dextran), which is too large for muLEC uptake (van Lessen et al, 2017) and serves as an injection efficiency control. Following ventricular injection, larvae were imaged by confocal microscopy for several hours to measure the increase of the pHrodo-avidin signal within muLECs (Figure 2a). In a representative larva sampled at high frequency (every 4 minutes), muLECs steadily accumulated pHrodo-avidin over the imaging window in a sigmoidal pattern (Figure 2b). To enable sufficient experimental throughput, we tested whether muLEC accumulation of pHrodo-avidin could be accurately modelled if down sampled (once every 60 minutes) and found that the data is well captured by a simple linear regression model with no apparent biases to early, middle, or late imaging starting points post-injection (Figure S1a). Therefore, in subsequent experiments we used the slope from a simple linear regression to quantify muLEC-uptake of pHrodo-avidin.

**Figure 2:**
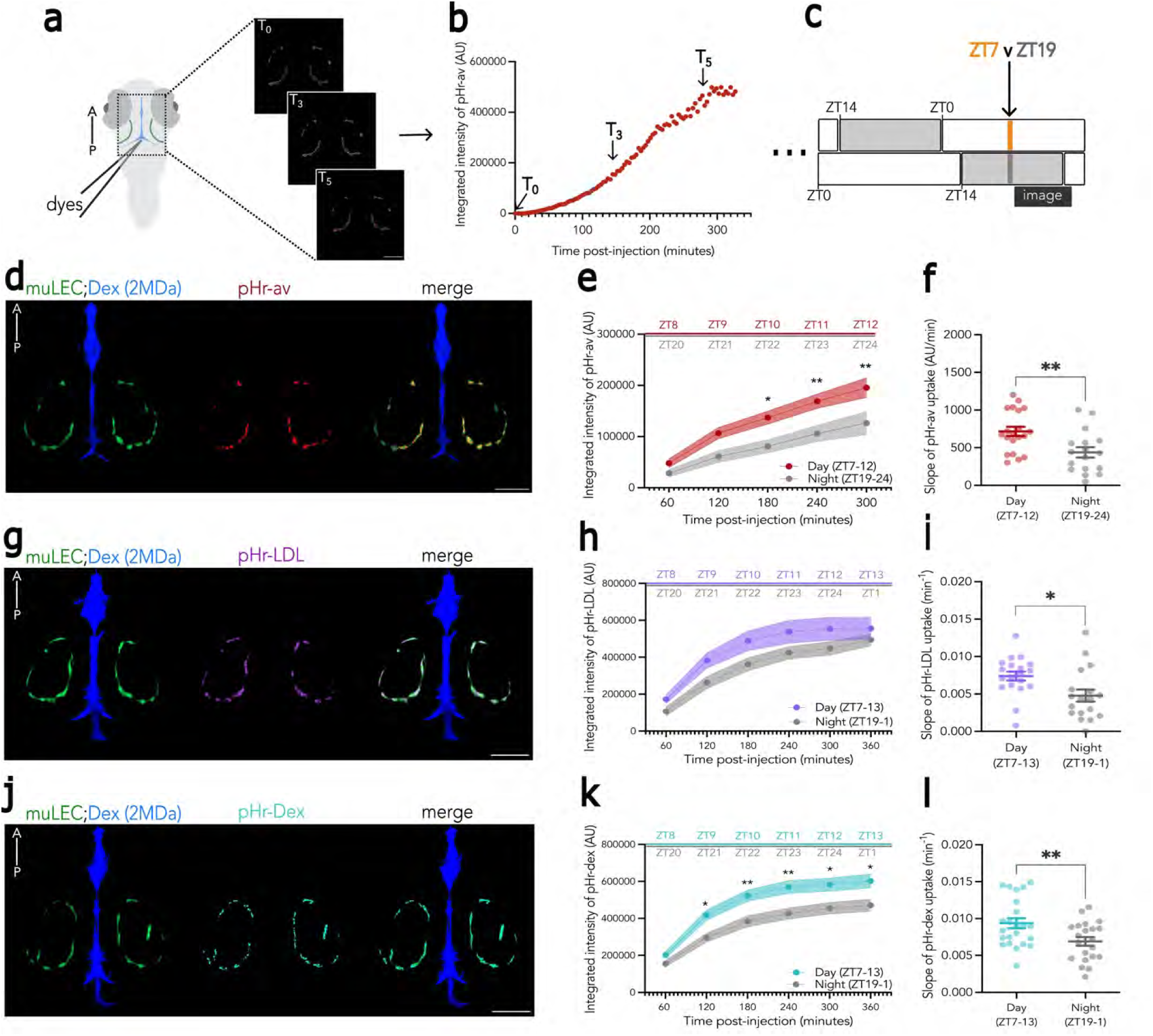
muLEC-uptake of macromolecules fluctuates over the day-night cycle. **a)** *Tg(flt4:mCitrine)* larva were intraventricularly injected with dyes and imaged for several hours to track muLEC macromolecular uptake. **b**) Time-course of pHrodo-avidin accumulation in muLECs, sampled every 4 minutes. **c**) Developmentally matched larvae on reverse LD cycles were injected and imaged in the middle of the day (ZT7) or night (ZT19). **d**) Representative maximum projection from a timeseries of a *Tg(flt4:mCitrine)* larva injected with 2MDa Alexa-fluor 488 dextran (Dex; blue) and pHrodo Red avidin (pHr-av; red). Signal was extracted from muLEC and ventricle masks removing any brain derived signal. **e**) The average integrated intensity ± SEM (60 min bins) of pHrodo-avidin uptake by muLECs is significantly higher in the day (ZT7-12, n=20) than night (ZT19-24, n=17). **f**) Slope of muLEC-uptake of pHrodo-avidin from a simple linear regression. **g**) Larva as in d) but injected with pHrodo Red LDL (pHr-LDL; magenta). **h, i**) muLEC uptake of pHrodo-LDL is higher during the day (ZT7-13, n=19) than at night (ZT19-1, n=18); the slope is from a one phase association regression. **j**) Larva as in d) but injected with pHrodo Red dextran (10kDa) (pHr-dex; cyan). **k**, **l**) muLEC-uptake of pHrodo-dextran is higher during the day (ZT7-13, n=23) than at night (ZT19-1, n=21). Data are mean ± SEM; each dot represents one larva (**f, i, l**). Statistical differences were determined by Mixed-Effects Model (**e, h, k**) or one-way ANOVA (two-tailed, **f, i, l**). ^ns^ p>0.05;* p< 0.05; ** p<0.01. Scale bar = 100µm.

Using the multiplexed muLEC-uptake assay, we then asked whether muLEC-uptake rates of pHrodo-avidin varied over the day-night cycle. Developmentally matched *Tg(flt4:mCitrine)* larvae raised on opposing light schedules were co-injected with 2MDa-dextran and pHrodo-avidin either in the middle of the day (ZT7, i.e., 7 hours after lights on) or night (ZT19, i.e., 5 hours after lights off) (Figure 2c, d). Larvae were imaged every 60 minutes for 5 hours post-injection with no difference in the integrated intensity of 2MDa-dextran at the first imaging time point, indicating similar injection volumes between groups (Figure S2e, p=0.95), and no time-sampling bias (Figure S2g, p=0.95). The rate of muLEC-uptake of pHrodo-avidin was higher during the late day compared to the middle of the night (Figure 2e,f; p=0.0044). This diurnal change in muLEC-uptake capacity of pHrodo-avidin was not observed when larvae were injected at the start of the day (ZT1) versus the start of the night (ZT14) (Figure S2b, c; p=0.42).

As muLECs take up a wide variety of macromolecules (Huisman et al, 2020), we next tested whether uptake of other cargo also varies across the day-night cycle. We co-injected larvae during the day or night (Figure 2c) with 2MDa-dextran and pHrodo conjugated to low-density lipoprotein (LDL; herein termed pHrodo-LDL) (Figure 2g) or pHrodo conjugated to 10kDa dextran (dex; herein termed pHrodo-dex) (Figure 2j) and imaged every 60 minutes for 6 hours. For both pHrodo-LDL (Figure S2h) and pHrodo-dex (Figure S2k), the rates of uptake were best captured at low sampling frequency by a one-phase association model. Similar to pHrodo-avidin (Figure 2e,f), the uptake rates for both pHrodo-LDL (Figure 2h,i; p=0.0135) and pHrodo-dex (Figure 2k,l; p=0.0089) were higher during the day compared to the night, with no differences in initial injection volume (pHrodo-LDL, Figure S2i, p=0.95; pHrodo-dex, Figure S2l, p=0.95) or in time-sampling (pHrodo-LDL, Figure S2j, p=0.95; pHrodo-dex, Figure S2m, p=0.95) between groups. Thus, muLECs are more capable of internalizing a variety of macromolecules during the waking day than at night.

### Uptake of pHrodo-avidin is regulated by a muLEC-autonomous circadian clock

The variation in muLEC uptake across the 24-hour day-night cycle could be regulated by three co-varying, yet experimentally separable factors: the direct effects of light exposure; the propensity to be awake (during the day) or asleep (at night); or the timing of the molecular circadian clock (Figure 3a). We therefore designed a series of pharmacological, genetic, and behavioural experiments to disentangle which of these factors are regulating the daytime increase in muLEC uptake rates, focusing on pHrodo-avidin.

**Figure 3:**
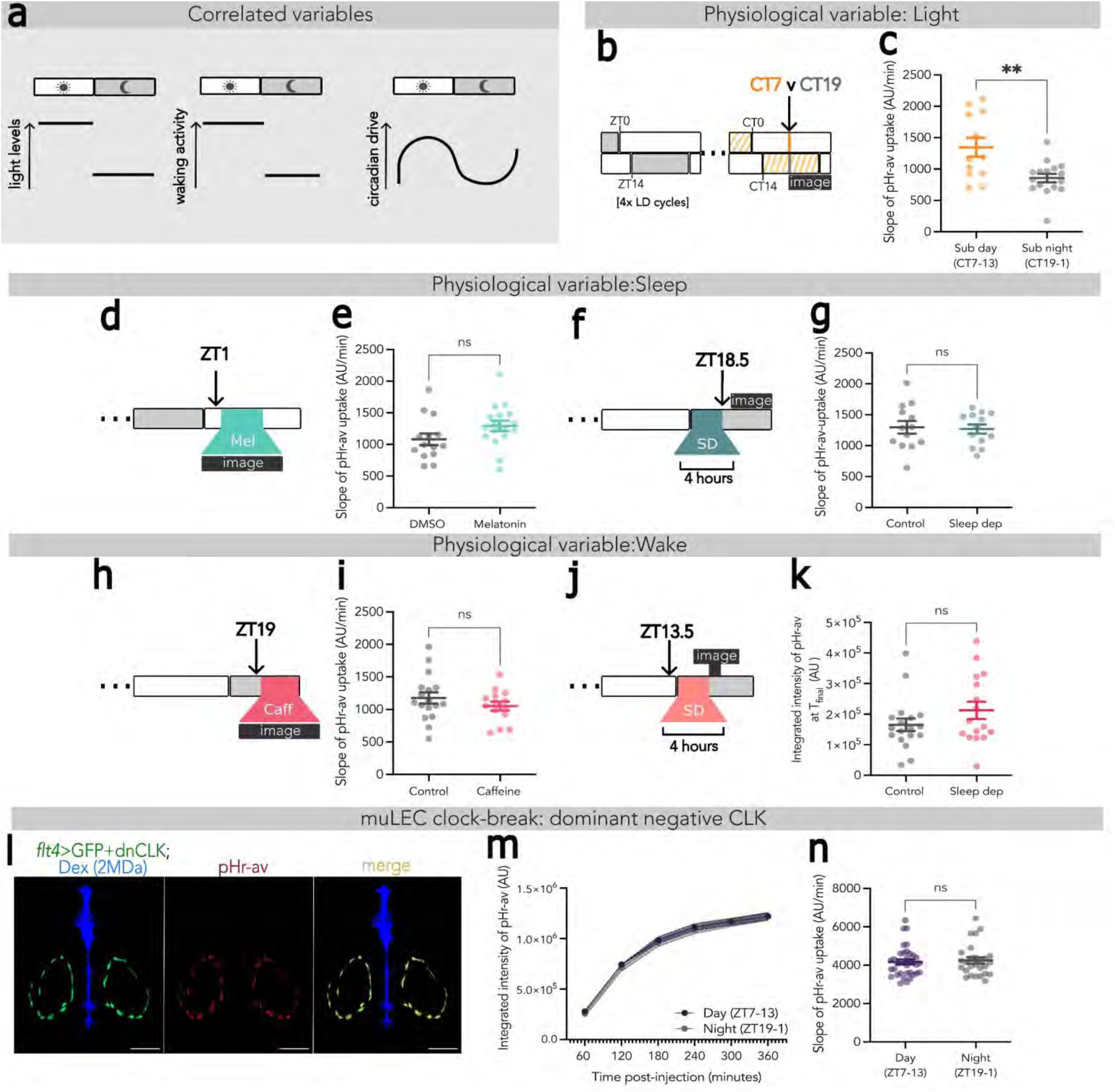
A muLEC-intrinsic circadian clock regulates the diurnal variation of pHrodo-avidin uptake. **a**) Candidate physiological drivers of uptake rates covary across the day-night cycle. **b**) muLEC-uptake rates were measured under free-running constant light conditions following injection (black arrow) during the subjective day (CT7) or night (CT19). **c**) The slope of muLEC uptake of pHrodo-avidin is higher during the subjective day (CT7-13, n=13) than subjective night (CT19-24, n=16). **d**) Experimental design to test the effect of melatonin. **e**) muLEC uptake of pHrodo-avidin is not significantly different between melatonin (n=18) treated and DMSO controls (n=19). **f**) Experimental design to test the effect of recovery sleep after sleep deprivation. **g**) muLEC uptake of pHrodo-avidin is not significantly affected by recovery sleep (n=14) vs controls (n=12). **h**) Experimental design to test the effect of caffeine. **i)** muLEC uptake of pHrodo-avidin is not affected by caffeine (n=15) vs controls (n=17). **j)** Experimental design to test the effect of “gentle handling” sleep deprivation. **k**) muLEC uptake of pHrodo-avidin during 4 hours of sleep deprivation (n=16) is similar to undisturbed controls (n=18). **l**) Timeseries maximum projection of a *Tg(flt4:Gal4); Tg(UAS:GFPp2AdnCLK)* larva injected with 2MDa Alexa-fluor, 488 dextran (Dex; blue) and pHrodo Red avidin (pHr-av; red). Signal was extracted from muLEC and ventricle masks removing any brain derived signal. **m**,**n)** The average integrated intensity ± SEM (m) and slope of muLEC-uptake of pHrodo-avidin by simple linear regression (n) are not significantly different between the day (n=35) and night (n=27). Data are mean ± SEM and each dot represents one larva (**c,e,g,i,k,n**). Statistical differences were determined by: one-way ANOVA (two-tailed, **c,e,g,j**). ^ns^ p>0.05;** p<0.01. Scale bar = 100µm.

First, to test whether the day-night difference in pHrodo-avidin uptake is directly influenced by light, we tracked muLEC uptake in larvae that were entrained on 14hr:10hr light-dark cycles to 4-5dpf then switched into constant light (free-running conditions) for 1 day. Although light levels are constant across the 24-hour day, the phases of both the circadian clock (Kaneko and Cahill, 2005) and the diurnal timing of wake and sleep states are preserved (Hurd and Cahill, 2002; Suppermpool et al, 2024). Developmentally matched *Tg(flt4:mCitrine)* larvae were then injected either during the middle of the subjective day (circadian time 7, CT7) or night (CT19) as entrained by the prior light-dark cycles (Figure 3b). Under these free-running conditions, muLEC-uptake of pHrodo-avidin remained higher during the subjective day than during the subjective night (Figure 3c, p=0.0035) with no difference in loading injection volume between groups (Figure S4a, p=0.1) or time sampling (Figure S4b, p=0.54). Since light exposure was the same at both timepoints, the subjective day increase in muLEC uptake cannot be due to signalling events that directly depend on a light-dark cycle alone.

Next, we assessed whether the 24-hour dynamic of pHrodo-avidin uptake is modulated by brain state by acutely modulating sleep and wake behaviours independently of the lighting conditions and time of day. To test whether acute inductions of sleep modulated the rate of pHrodo-avidin uptake, larvae were injected in the morning (ZT1) when locomotor activity is typically high and then transferred into a solution containing either 1µM melatonin dissolved in 0.2% dimethyl sulfoxide (DMSO) or a 0.2% DMSO control (Figure 3d). In zebrafish, melatonin is an acute hypnotic that works downstream of the circadian clock to regulate sleep (Zhdanova et al, 2001; Gandhi et al, 2015); melatonin-treated fish will therefore have a higher propensity to sleep while remaining circadian phase-matched to their DMSO-treated conspecifics (Suppermpool et al, 2024). muLEC-uptake of pHrodo-avidin remained the same between melatonin and DMSO treated larvae (Figure 3e, p=0.098). In a second sleep assay, we acutely but temporarily exposed larvae to the GABA-A receptor antagonist, pentylenetetrazol (PTZ, 10mM), which induces a prolonged increase in rebound sleep during the day with no effect on molecular circadian rhythms (Reichert et al, 2019; Benoit et al, 2024), and then assessed muLEC uptake capacity for pHrodo-avidin (Figure S3a). Similar to melatonin-treated fish, post-PTZ-induced daytime sleep did not impact muLEC-uptake rates compared to fish-water treated controls (Figure S3b, p=0.84).

Because inducing sleep during the day with drugs may not fully recapitulate natural night-time sleep states, we also assessed whether promoting even more sleep during the subjective night could impact muLEC rate measurements. To do this, we used a gentle handling sleep deprivation protocol to keep larvae awake for 4hr at the start of the night (ZT14-ZT18), which is followed by increased sleep consolidation and duration (Suppermpool et al, 2024; Elias et al, 2025). We then injected pHrodo-avidin immediately after sleep deprivation and tracked muLEC uptake for the remainder of the night period (ZT19-24) (Figure 3f). Again, we observed no change in uptake rates in this post-sleep deprivation period (Figure 3g, p=0.82). Taken together, these results indicate that muLEC uptake of pHrodo-avidin is not modulated by acute inductions of sleep, either during the day or night, suggesting that the lower rate of uptake during the subjective night cannot be due to increased sleep alone.

We then tested whether uptake of pHrodo-avidin could instead be boosted by increased wakefulness at night. Larvae were injected in the middle of the night (ZT19), immediately transferred into a solution containing 250µM caffeine to promote wakefulness, and uptake was tracked for the remainder of the night (Figure 3h). This caffeine-induced wakefulness did not affect muLEC uptake rates (Figure 3l, p=0.28). To test the effects of natural waking states on uptake, larvae were injected with pHrodo-avidin prior to the start (ZT13.5) of 4hr of gentle handling sleep deprivation, and muLEC uptake rates were imaged immediately afterward at ZT18 (Figure 3j). Again, no difference in uptake of pHrodo-avidin was observed during sleep deprivation (Figure 3k, p=0.35), demonstrating that boosting wakefulness with either drugs or gentle handling does not alter the muLEC’s capacity to internalize pHrodo-avidin.

Since neither acute light exposure nor sleep-wake state can explain the diurnal variation in pHrodo-avidin uptake, we next examined whether the difference could be due to non-behavioral outputs of the circadian clock. Zebrafish embryos raised from fertilization under constant light and temperature conditions fail to develop coherent 24-hour circadian rhythms, as assessed by the loss of rhythmic clock gene expression and behaviour (Whitmore et al, 2000; Prober et al, 2006; Wang et al, 2020; Suppermpool et al, 2024). We therefore tested the muLEC uptake of pHrodo-avidin in larvae under these continuous light “clock-break” conditions compared to developmentally matched larvae under entrained, free-running conditions (Figure S3c). Larvae under free-running conditions had higher levels of pHrodo-avidin accumulating in muLECs relative to “clock-broken” larvae when sampled in the subjective day (CT7) but not during the subjective night (CT19) (Figure S3d; clock-break vs CT7-13, p=0.0125; clock-break vs CT19-1 p=0.346). We conclude that day-night differences in pHrodo-avidin uptake rely on intact circadian rhythms.

Zebrafish larvae, like many species, have endogenous molecular circadian oscillators in most, if not all, cells (Whitmore et al, 2000; Mohawk et al, 2012; Moore & Whitmore, 2014). Therefore, the clock-dependent diurnal variation in pHrodo-avidin uptake could be due either to extrinsic circadian cues that signal onto muLECs (non-cell autonomous), or to muLEC-intrinsic circadian rhythms (cell-autonomous) that boost muLEC scavenger function during the subjective day. To test this, we expressed in muLECs a dominant negative Clock (dnCLK), which can bind to its partner BMAL but lacks the ability to bind DNA and direct transcription of key circadian clock genes, thereby cell-autonomously disrupting the essential positive arm of the transcription-translation feedback loop that underlies intrinsic circadian timekeeping (Dekens & Whitmore, 2008; Livne et al, 2016; Kelu et al, 2020). Double transgenic *Tg(flt4:Gal4);Tg(UAS:GFPp2AdnCLK)* larvae showed robust and uniform expression of the dnCLK construct in muLECs at 5/6 dpf (Figure 3l), and this dnCLK expression abolished the day-night difference in pHrodo-avidin uptake (Figure 3m, n; p=0.7). Thus, a muLEC-intrinsic circadian oscillator regulates the daily rhythm of pHrodo-avidin uptake capacity.

### The muLEC-intrinsic clock regulates day-night uptake through the rhythmic expression of Mrc1a protein

An attractive hypothesis for how a muLEC-intrinsic circadian clock might dictate day-night uptake rates of pHrodo-avidin is through timed regulation of the molecular machinery required for uptake, via transcriptional or post-transcriptional mechanisms. We therefore focused on the pattern-recognition receptor Mannose receptor C-type lectin 1a (Mrc1a), which had previously been implicated in mediating uptake in muLECs (van Lessen et al, 2017; Huisman et al, 2021). First, we confirmed the requirement of *mrc1a* for uptake of avidin by injecting *Tg(flt4:mCitrine)* embryos at the one-cell stage with Cas9 and guideRNAs targeting three separate *mrc1a* exons to generate *mrc1a* F_0_ knockouts (KO) (Figure 4a; S5a-d). We confirmed that *mrc1a* KOs harboured a high-proportion of frameshift mutations (Figure S5e) upstream of sequences encoding key Mrc1a protein domains (Figure S5b), strongly indicating that *mrc1a* KOs are loss of function mutants. Unexpectedly, compared to sibling controls injected with non-targeting gRNAs, *mrc1a* KOs exhibited abnormal muLEC morphology (Figure 4b; S6a), with an increase in total cell volume (Figure S6b) and mislocalisation of muLECs inside the tectal rings, an ectopic distribution for this stage of development (Figure S6c), even though there were no differences in the surrounding blood vasculature (Figure S6d-f) or microglia (Figure S6g-k). Mrc1a must therefore play a key role in regulating either the differentiation, cell division, or migration of muLECs at this stage as has been previously described in macrophages (Sturge et al, 2007). Nevertheless, despite the larger muLEC pool, and in line with previous work (Huisman et al, 2021), *mrc1a* loss-of-function completely abolished muLEC uptake of pHrodo–avidin at all measured time points post-injection (Figure 4c; 4d, p<0.0001). Interestingly, round and dynamic pHrodo-avidin signal was still detected in the brain in *mrc1a* KOs suggesting that Mrc1a is not required for pHrodo-avidin uptake by brain resident phagocytes (Figure 4b). This indicates that Mrc1a functions as a non-redundant receptor for avidin internalization and muLECs have no compensatory pathways.

**Figure 4:**
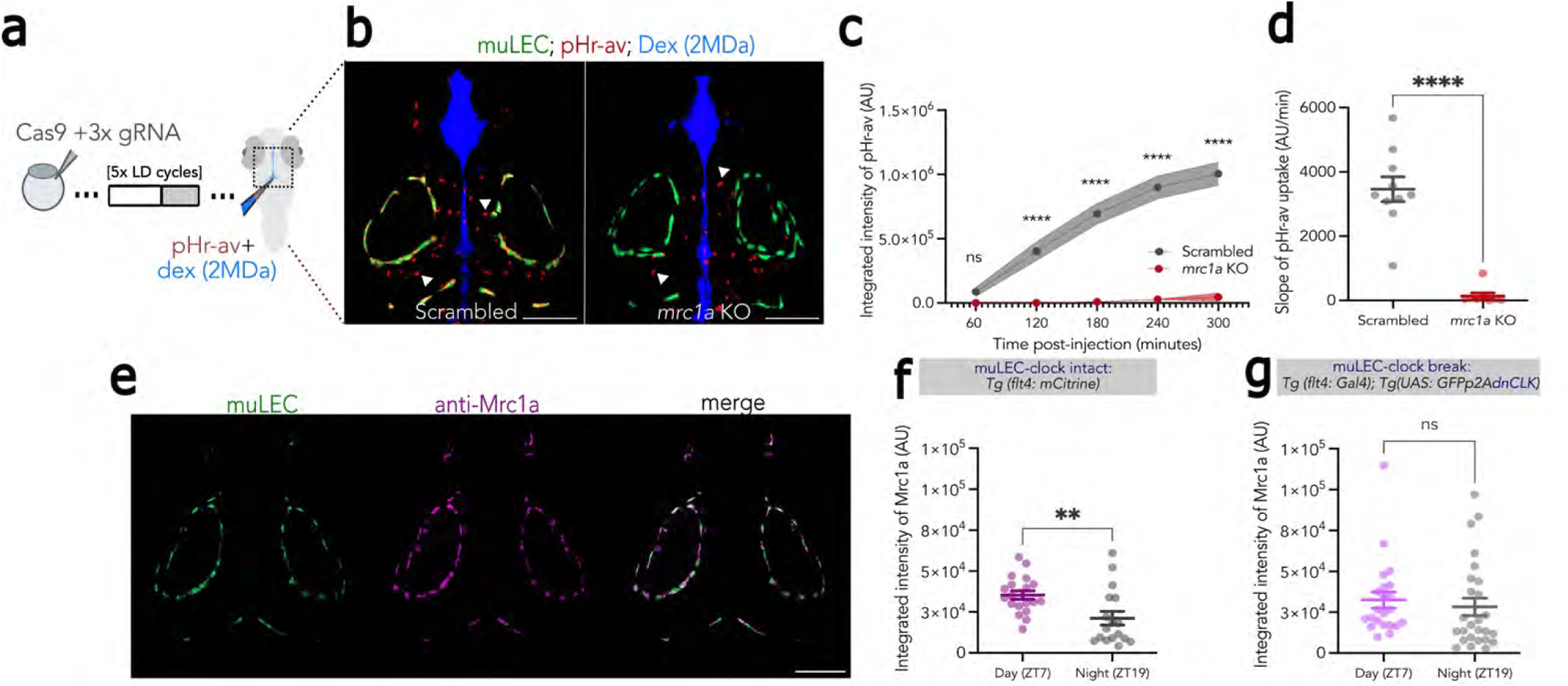
muLEC-intrinsic circadian regulation of Mrc1a protein regulates pHrodo-avidin uptake. **a**) The experimental paradigm. *Tg(flt4:mCitrine)* larvae were injected at the one-cell stage with Cas9 and *mrc1a*-targeting or scrambled control gRNAs, then injected with dyes at 5dpf. **b**) Representative maximum projection from a timeseries of a scrambled control larva (left) and *mrc1a* KO (right) with muLECs (green) post-intraventricular injection of 2MDa Alexa-fluor, 488 dextran (Dex; blue) and pHrodo Red avidin (pHr-av; red). White arrowheads indicate pHr-avidin uptake by brain-resident phagocytes. **c**) The average integrated intensity ± SEM of pHrodo-avidin accumulation in *mrc1a* KO larvae (n=8) and scrambled control siblings (n=10). **d**) The slope of muLEC-uptake of pHrodo-avidin. **e**) Representative maximum projection of muLECs (green) and anti-Mrc1a (magenta) immunofluorescence in 5dpf *Tg(flt4:mCitrine)* larvae. **f**) Integrated intensity of anti-Mrc1a immunofluorescence in muLECs is higher during the day (ZT7, n=19) than night (ZT19, n=17). **g**) Integrated intensity of anti-Mrc1a immunofluorescence in muLECs is similar in the day (n=22) and night (n=26) in muLEC-clock disrupted larvae. Data are mean ± SEM. Each dot represents one larva (c,d,f,g). Statistical differences were determined by a Mixed-Effects Model (c) or Mann-Whitney U tests (two-tailed, d,f,g). ). ^ns^ p>0.05; ** p<0.01; ****p<0.0001. Scale bar = 100µm.

We then examined whether Mrc1a expression or availability changes in a circadian clock-controlled manner by performing immunohistochemistry during the day or night with a custom anti-Mrc1a antibody (Figure 4e; S5f) on muLEC-clock-intact (*Tg(flt4:mCitrine))* or muLEC-clock-broken (*Tg(flt4:Gal4; Tg(UAS:GFPp2AdnCLK))* larvae. In muLEC-clock intact larvae, Mrc1a protein expression was significantly higher in the day compared to the night (Figure 4f, p=0.0048), but this diurnal variation was abolished in muLEC-clock broken larvae (Figure 4g, p=0.135). These results are consistent with a model in which the muLEC-intrinsic circadian clock drives higher daytime Mrc1a protein levels that enhance muLEC capacity for the internalization of pHrodo-avidin.

### Diurnal variation of LDL uptake is regulated by sleep-wake state and not a muLEC-intrinsic clock

Unlike pHrodo-avidin, muLEC-uptake of both pHrodo–dex and pHrodo-LDL occur in an Mrc1a-independent manner (Huisman et al, 2021). However, despite relying on different receptor machinery, is the diurnal uptake of carbohydrate and lipoprotein similarly controlled by a muLEC-autonomous circadian clock? To test this, larvae expressing dnCLK in muLECs were injected with either pHrodo-dex or pHrodo-LDL in the day and night, and the muLEC accumulation of these substrates was tracked for several hours (Figure 5a). Despite interference with the muLEC-intrinsic clock, uptake of both pHrodo-dex (Figure 5b, p=0.0039; Figure S7a) and pHrodo-LDL (Figure 5c; p=0.03; Figure S7d) remained higher during the day than night, showing that the muLEC-intrinsic clock is insufficient to drive diurnal changes in the uptake of LDL and dextran.

**Figure 5:**
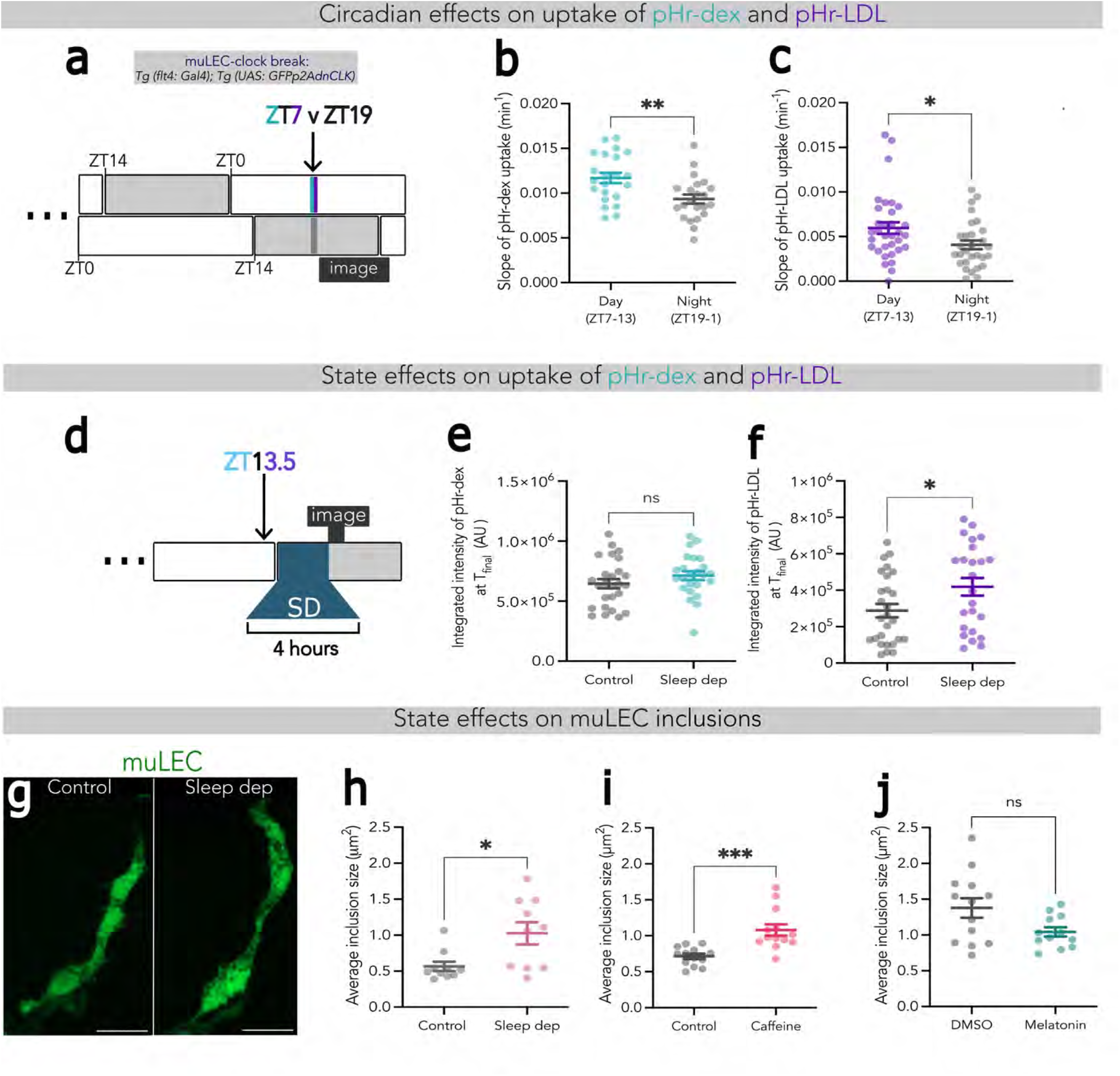
Diurnal variation in pHrodo-dex and pHrodo-LDL uptake are independent of the intrinsic clock. **a**) The experimental paradigm to measure day (ZT7) vs. night (ZT19) muLEC uptake rates in *Tg(flt4:Gal4); Tg(UAS:GFPp2AdnCLK)* larvae. **b,c**) Slopes of muLEC-uptake of pHrodo-dex (b) pHrodo-LDL (c) from a one-phase association regression show that uptake differences between day (pHr-dex, n=22; pHr-LDL, n=33) versus night (pHr-dex, n=22; pHr-LDL, n=29) persist despite transgenic muLEC-clock disruption. **d**) Experimental design to test the effect of extended waking on muLEC uptake of pHrodo-dex and pHrodo-LDL. **e**) The integrated intensity of pHrodo-dex accumulation is similar during 4 hours of sleep deprivation (n=24) vs controls (n=26). **f**) The integrated intensity of pHrodo-LDL accumulation is significantly higher after sleep deprivation (n=24) vs controls (n=27). **g**) A muLEC from a *Tg(flt4:mCitrine)* larva following 4 hours of sleep deprivation (right) or undisturbed control (left). **h**) Average size of intracellular muLEC inclusions after sleep deprivation (n=10) vs controls (n=10). **i**) Average size of muLEC intracellular inclusions at ZT23 after 4 h of Caffeine exposure (250μM, n=13) or water control (n=13). **j**) Average size of muLEC intracellular inclusions at ZT5 after 4 h of Melatonin exposure (1μM, n=13) or DMSO control (n=13). Data are mean ± SEM; each dot represents one larva. Statistical differences were determined by Mann-Whitney U tests (two-tailed, c,f,h,j) or one-way ANOVA (two-tailed, b,e,i). ^ns^ p>0.05;* p<0.05; ** p<0.01; p<0.001. Scale bar:10µm.

We then asked whether pHrodo-dex and pHrodo-LDL uptake are instead governed by non-circadian processes, such as brain state (e.g., sleep vs. wake). To test this, *Tg(flt4:mCitrine)* larvae were injected with pHrodo-dex or pHrodo-LDL before the start of the night (ZT13.5) and either kept awake by gentle handling for four hours (ZT14-18) or left to sleep undisturbed (Figure 5d). Similar to pHrodo-avidin uptake (Figure 3j), forced wakefulness did not impact muLEC-uptake rates of pHrodo-dex (Figure 5e, p=0.216). Intriguingly, however, sleep deprivation significantly increased the muLEC-uptake rates of pHrodo-LDL compared to undisturbed controls (Figure 5f, p=0.035). Furthermore, inducing night-time wakefulness either by sleep deprivation or 250µM caffeine treatment significantly increased the size (Figure 5h, p=0.0012; Figure S7k; Figure 5i p=0.0003; Figure S7o) but not the number of muLEC inclusions (Figure S7h, p=0.156; S7l, p=0.13), while forced daytime sleep with melatonin trended towards a decrease in inclusion size (Figure 5j, p=0.098; Figure S7s) with no change in the average number of inclusions (Figure S7j, p>0.99) or filopodia measurements (Figure S7i,j ; S7m,n ; S7q,r). Since LDL is trafficked into larger, acidified late endosomes (Islam et al, 2022), these observations are consistent with the heightened increase in LDL uptake seen after extended wakefulness.

Thus, although muLECs have a higher capacity for all three macromolecular cargoes during the waking day, this increase is accomplished by at least two, and possibly three, independent mechanisms: avidin relies on a muLEC-autonomous clock; LDL is attuned to the waking state of the brain; and dextran requires neither.

### muLECs facilitate behavioural homeostasis post-seizure

Taken together, muLEC scavenging is modulated by circadian and behavioural state to peak during the day when metabolic demand is greatest. Consequently, we hypothesised that muLECs contribute to maintaining brain solute homeostasis during metabolic challenges and that their disruption would impact physiological recovery following a metabolically taxing insult. To test this, we used short multiphoton laser pulses to ablate muLECs, which eliminated all muLECs while leaving the surrounding blood vasculature intact (Figure 6a). Mock ablations were performed by aiming the laser adjacent to muLECs. The mock- and muLEC-ablated larvae were then treated with 10mM PTZ for 1 hour to induce seizures (Figure S3a) and subsequently video-tracked for 48 hours in free-running conditions. Representative traces show that both mock- and muLEC-ablated larvae have an initial multi-hour period of extreme inactivity following PTZ treatment, but all larvae eventually resume normal levels of locomotor activity (Figure 6b), with no difference in daytime (Figure 6d, p=0.329) or night-time activity levels between mock and ablated groups after recovery (Figure 6e, p=0.592). However, muLEC-ablated larvae took significantly longer (p<0.05, rank sum test) to recover normal locomotor activity (defined by reaching activity levels equal to their next days’ 50^th^ percentile) than mock-ablated larvae (Figure 6c), with 50% of muLEC-ablated larvae recovering after 11.38 hours compared to 7.75 hours for mock-ablated larvae. This delay in post-seizure recovery was consistently found irrespective of the percentile cut-off used to define recovery (Figure S8a). These data suggest that muLECs contribute to re-establishing solute homeostasis after a metabolically taxing insult, which may also explain why muLEC function is boosted to meet the higher metabolic demands of the waking day.

**Figure 6.**
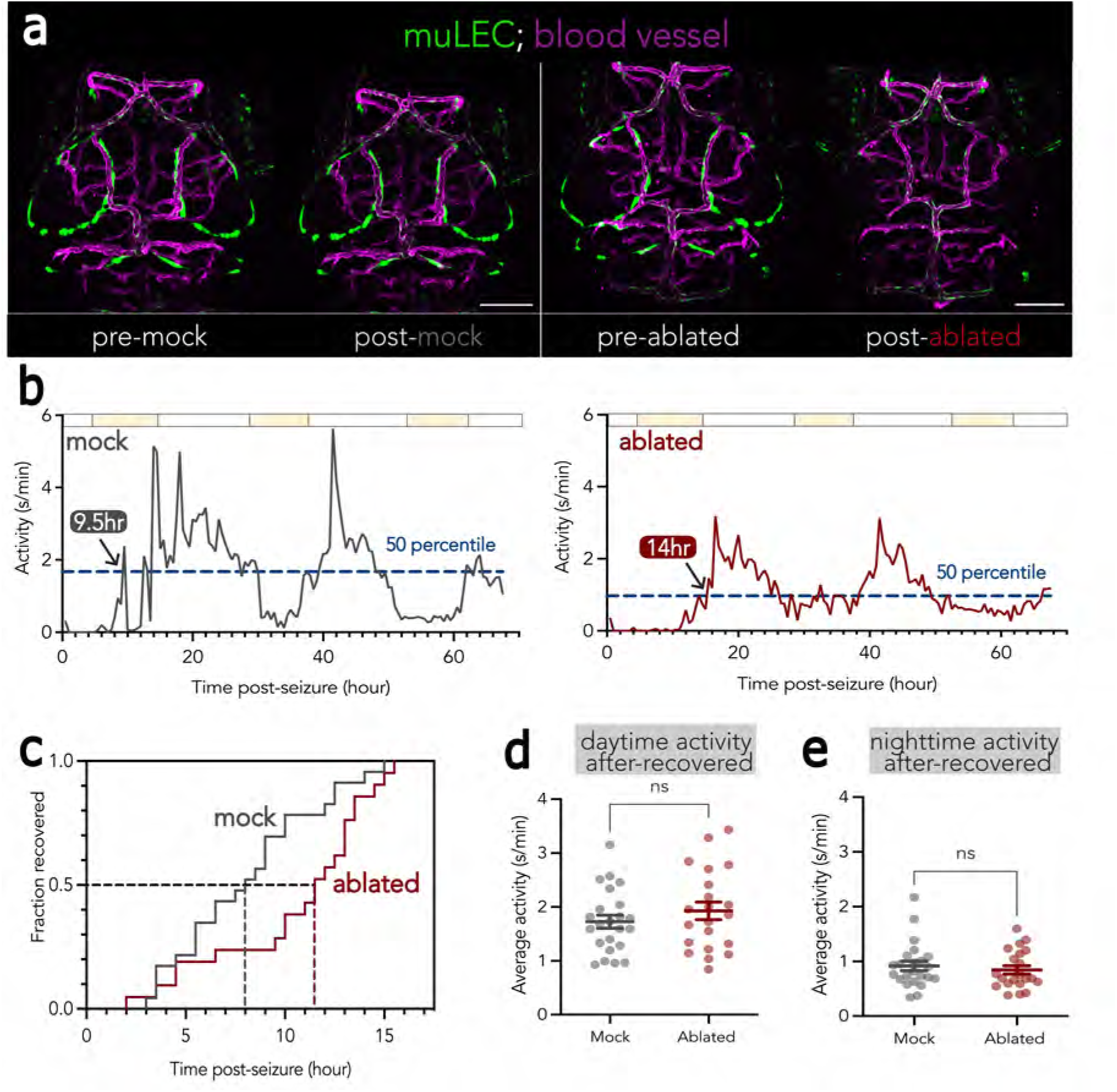
muLECs facilitate behavioural recovery following drug-induced seizures. **a)** Maximum projections of confocal stacks pre- and post-mock (left) or pre- and post-targeted laser ablation (right) of muLECs (green) in a 5dpf *Tg(flt4:mCitrine);Tg(kdrl:mCherry)* larva. Blood vessels (magenta) are unaffected. **b**) Multi-day activity traces showing the recovery of baseline behavior following PTZ-induced seizures (1hr ending at time 0). A representative larva from mock- (left, grey) and muLEC-ablated (right, red) conditions are shown. Recovery is determined by the time the activity levels reach the 50th percentile of each larva’s post-recovery baseline activity level (dashed navy line). **c**) The fraction of fish that have recovered by a given time post seizure. Mock ablated fish recover faster than muLEC-ablated fish (log rank test, p<0.05). **d, e**) Average daytime (**d**) and nighttime (**e**) activity of ablated (n=21) and mock ablated (n=23) fish is similar after recovery. Statistical differences were determined by Mann-Whitney U tests (two-tailed, e); one-way ANOVA (two-tailed, d). ^ns^ p>0.05; *** p<0.001. Scale bar:100µm.

## Discussion

Here, we demonstrate that at least two distinct mechanisms converge to boost CSF solute clearance by a meningeal scavenger cell population during the waking day and that this function is critical for recovery after a brain metabolic challenge. Given that the muLECs are poised to participate in maintaining solute homeostasis of the CSF, these results have important implications for the regulation and timing of brain clearance.

The brain is metabolically demanding during the waking day, and this is well reflected in the content of the CSF. The total protein content of the CSF undergoes diurnal fluctuation and is higher in the latter portions of the waking day (Kakarla et al, 2023). The CSF abundance of a range of solutes spanning the neuromodulators dopamine (Poceta et al, 2009), cortisol (Kalin et al, 1980), and hypocretin (Zeitzer et al, 2003) as well as the waste products amyloid β (Bateman et al, 2007; Huang et al, 2012; Lucey et al, 2015) and tau (Holth et al, 2019) all peak during active periods as a function of brain state or circadian cues. Additionally, peroxidated lipids build up during the waking day and accumulate in glia as lipid droplets (Haynes et al, 2024), while sleep disruption has been shown to alter the brain (Rorsman et al, 2025) and CSF lipidome (Koethe et al, 2009; Saito et al, 2021; Dakterzada et al, 2023) as well as the CSF distribution of lipid-associated apolipoproteins (Achariyar et al, 2016). Together, these findings demonstrate that routes exist to connect the ISF of the brain to the CSF within the surrounding meninges and further argue that wakefulness is a period of active solute release into the CSF that creates a demand for the removal of an array of macromolecular waste during behaviourally active periods.

Our data show that muLEC-uptake of proteins, lipoproteins, and carbohydrates are all higher during the day, with uptake of specific proteins relying on the timed expression of Mrc1a via a cell-autonomous clock. In contrast, lipoprotein uptake is attuned to, and regulated by, wake-dependent cues, independently of the circadian clock. Why use two distinct mechanisms to regulate protein versus lipoprotein uptake, given that under normal conditions, both lead to a co-ordinated increase during the waking day? One hypothesis is that each uptake pathway is regulated in accordance with the clearance demands of the endogenous substrate/s that it serves. Since the circadian clock anticipates the 24-hour day but does not typically allow for more dynamic modulation, it could be that this cell-autonomous regulation serves to clear predictably oscillating Mrc1a-dependent substrates from the CSF such as glycosylated endo-lysosomal proteins (van der Zande et al, 2021; Cummings, 2022; Orduña Dolado et al, 2024). Alternatively, muLECs might provide an important barrier to prevent proteinaceous material entering the CSF/brain; such risks might be more reliably likely to occur during active behavioural exploration. In contrast, the regulation of lipoprotein uptake may require faster kinetics in response to more acute events like sleep deprivation, when harmful material such as oxidized lipids start to accumulate (Haynes et al., 2024). An open question for future work is to what extent microglia and other endocytic cell populations also engage in this dual-mode of regulation.

Another important question remains as to which wake-related signal(s) muLECs are attuned, and how this signal(/s) is translated within muLECs to potentiate LDL uptake. A recent study reported that muLEC development is dependent on, and regulated by, neuronal activity via stimulation of radial glia that release lymphangiogenic vascular endothelial growth factor C and D (Vegfc, Vegfd) into the meninges (Li et al, 2025). It is not yet clear whether this neuron glia muLEC signalling cascade is limited to controlling only muLEC development or could also be used to link brain state to the muLEC scavenging functions that we have tested here. The release of other VEGF signals has been shown to upregulate expression of the lipid scavenger receptors Stabillin-1 and low-density lipoprotein receptor-related protein 1 (LRP1) and to mediate lipid uptake and metabolism by macrophages and liver sinusoidal cells in the periphery (Wary et al, 2003; Hagberg et al, 2010; Falkevall et al, 2017; Shew et al, 2018; Tirronen et al, 2018; Ning et al, 2020; Luo et al, 2022). Future work should examine whether lipid uptake by muLECs is also regulated by VEGF-signalling and whether this provides a mechanism to link active brain states to muLEC function.

Finally, we demonstrated that muLECs are required to restore behavioural homeostasis following drug-induced seizure. It is well documented that seizures alter the composition of CSF, leading to elevated total protein content, lactate, and tau (Chatzikonstantinou et al, 2015; Tumani et al, 2015; Süße et al, 2019; Langenbruch et al, 2021). Additionally, there are elevated CSF levels of somnogenic cytokines such as interleukin-6 (Peltola et al, 1998; Vgontzas et al, 1999; Peltola et al, 2000; Rohleder et al, 2012) and meningeal-derived prostaglandins that promote post-ictal sleep and recovery (Kaushik et al, 2014; Mello & Oliveira, 2014). Therefore, it is possible that muLEC scavenging is critical for refining the CSF milieu in response to changing brain states, especially under insult. Future work should explore whether CSF solute clearance by muLECs contributes to other disease states known to impact CSF composition, such as in traumatic brain injury, chronic sleep disorders, CNS infections, or neurodegenerative disease (Morganti-Kossman et al, 1997; Vgontzas et al, 1997; Prasad et al, 2014; Carvalho et al, 2022).

To conclude, we demonstrate that CSF solute clearance by muLECs is differentially regulated by state and circadian cues to peak during the waking day. We propose that muLECs represent one node of many mechanisms involved in brain clearance and homeostasis. Understanding how the wake-active muLECs operate alongside other avenues of brain clearance proposed to undergo diurnal regulation will be important for a fuller picture of how brain waste is regulated in both normal and diseased brains.

## Acknowledgements

We thank Jeff Kelu and Simon Hughes for the UAS-dnCLK construct, Sandip Patel for consultation on pH-sensitive dyes, the Payne lab for MiSeq, the UCL Fish Facility for animal husbandry, and all members of the Rihel lab for constructive feedback on all stages of the project. The work was supported by a Wellcome Trust Investigator Award to JR (217150/Z/19/Z), a BBSRC grant to JR and TAH (BB/T001844/1), and a Medical Research Council Doctoral Training Partnership (MRC-DTP) fellowship awarded to TG.

## Author Contributions

TG, TK, JR and TAH conceived of the project. TK, with help from DK, imaged and analysed muLEC morphology in Figure 1 and 5g. TG designed, performed and analysed all uptake experiments (Figure 2-5) with help from LSM for Mrc1a immunostaining (Figure 4e-g). TG, TK, and JR performed muLEC ablation experiments (Figure 6). TG designed the figures and wrote the manuscript with input from all authors. JR supervised the project.

## Materials and Methods

### Animals

Zebrafish (Danio rerio) husbandry and experiments were conducted according to UCL Fish Facility standard protocols and under the project licenses PA8D4D0E5 and PP6325955 awarded to JR under the auspices of the UK Animal Scientific Procedures Act (1986). Embryos were kept in 10cm Petri dishes in E3 media (0.3 g/L Instant Ocean and 0.1% Methylene blue) in 14 hr-10 hr light–dark cycle incubators at 28 °C, except of clock break experiments, when they were kept under constant light from 0 dpf. Unless otherwise stated, all experiments were conducted on larvae 5-6 days post fertilisation. At the end of all experiments, larvae were euthanised using by anaesthetic overdose of MS222 (Sigma Aldrich, UK) or 2-phenoxyethanol (ACROS Organics).

All transgenic lines were kept in a *nacre (mitfa)^w2/w2^* mutant background. The following lines were used for experiments:

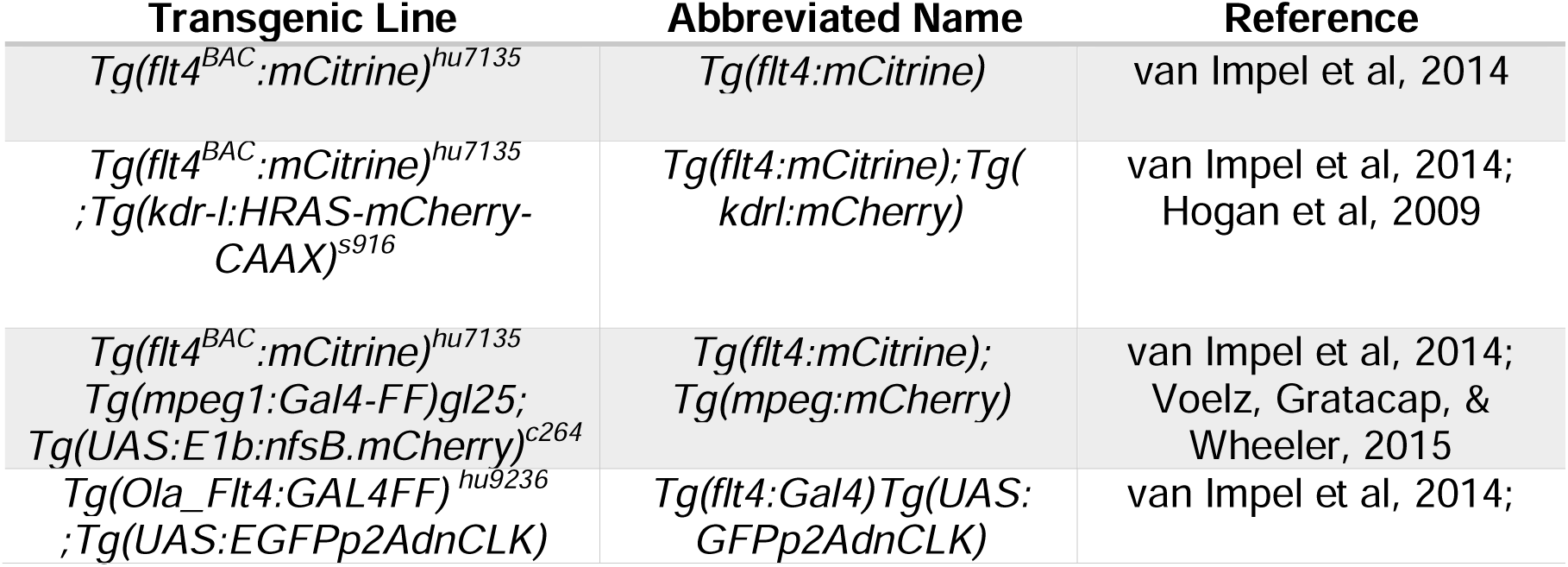

| Transgenic Line | Abbreviated Name | Reference |
| --- | --- | --- |
| *Tg(flt4<sup>BAC</sup>:mCitrine)*<sup>hu7135</sup> | *Tg(flt4:mCitrine)* | van Impel et al, 2014 |
| *Tg(flt4<sup>BAC</sup>:mCitrine)*<sup>hu7135</sup>;<br>*Tg(kdr-l:HRAS-mCherry-CAAX)*<sup>s916</sup> | *Tg(flt4:mCitrine);Tg(kdrl:mCherry)* | van Impel et al, 2014;<br>Hogan et al, 2009 |
| *Tg(flt4<sup>BAC</sup>:mCitrine)*<sup>hu7135</sup>;<br>*Tg(mpeg1:Gal4-FF)gl25*;<br>*Tg(UAS:E1b:nfsB.mCherry)*<sup>c264</sup> | *Tg(flt4:mCitrine);Tg(mpeg:mCherry)* | van Impel et al, 2014;<br>Voelz, Gratacap, & Wheeler, 2015 |
| *Tg(Ola_Flt4:GAL4FF)*<sup>hu9236</sup>;<br>*Tg(UAS:EGFPp2AdnCLK)* | *Tg(flt4:Gal4)Tg(UAS:GFPp2AdnCLK)* | van Impel et al, 2014; |

#### Transgenesis

A plasmid encoding the pT2-UAS-EGFP-p2A-dnCLK-5xMyc construct was kindly provided by Jeff Kulu and Yoav Gothlif. Approximately 1nl of plasmid (20ng/μl) alongside Tol2 transposase mRNA (50ng/μl) was injected in *Tg(flt4:Gal4FF)^hu9236^* (van Impel et al., 2014) embryos at the one-cell stage as previously described (Clark et al., 2011). F_0_ larvae from a single clutch were screened at 4dpf for positive EGFP expression and raised to adulthood. Founder adults were outcrossed to *nacre (mitfa)^w2/w2^* larvae to screen for transgene germline integration. A Nikon fluorescent SMZ1270i fluorescent stereo microscope was used to identify F_1_ progeny with the brightest EGFP fluorescence and further propagated. Experiments were performed on F_1_ or F_2_ larvae from a single parental clutch incross.

### muLEC cellular and subcellular morphology

To visualize and quantify muLEC morphology, 6dpf *Tg(flt4:mCitrine)* larvae were culled by anesthetic overdose (MS222, approximately 420 mg/L) and fixed in 4% paraformaldehyde (PFA). Following washing in 1x phosphate buffered saline (PBS) and 1x PBS with Triton (PBST), larvae were ventrally mounted in 1% low-melt agarose and imaged using a LSM 980 inverted confocal microscope with Airyscan 2 (Zeiss). For assessing muLEC volume, surface area, disconnectedness and width/length of muLEC connections a x20/1.0 NA objective was used and z-stacks were obtained at 1μm intervals of 4076 x 4084 px or 4060 x 4060 px and 16-bit using SR-8Y acquisition mode. For assessing muLEC inclusions and filopodia a x40/1.2 NA objective was used and z-stacks were obtained at 0.21 μm intervals of 4084 x 4084 px and 16-bit using CO-8Y acquisition mode. mCitrine was excited at 488nm.

Analysis of muLEC volume, surface area, disconnectedness were performed in Imaris. ROIs were constructed around all muLECs within one hemisphere and raw z-stacks were segmented and auto-thresholded to generate 3D surface reconstructions using the Imaris surface function. For quantification of the width/length of muLEC connections, 2D maximum projections were analysed in ImageJ (NIH) using a modified macro script from McDowell et al, 2020. For assessing muLEC inclusion sizes, 2D maximum projections were analysed in ImageJ with a custom-built macro (https://github.com/JRihel/muLEC-Circadian-Imaging). Filopodial lengths and numbers were manually analyzed using the segmented line tool in ImageJ.

### Intraventricular injection assay for assessing muLEC uptake

#### Lighting schedules

Larvae were reared in incubators kept at 28°C on various lighting schedules as specified (hours light: hours dark): light:dark, 14hr:10hr; free-running, 14:hr:10hr up to 4dpf then transferred to constant light; clock-break, 24hr:0hr, constant light. When multiple ZT times or different light schedules were compared within an experiment, water bath incubators were set up to run in parallel. Temperature and light transitions of water bath incubators were recorded with a HOBO Pendant (Onset Data Loggers) immersed in the water bath. If larvae were collected in their dark period, larvae were removed from the water baths using an external red head light (Blackburn Local Bike Rear Light 15 Lumen, UK) and transported in light proof boxes to be taken through the intraventricular injection assay where ambient light exposure was limited as much as possible.

#### Drug treatment

For PTZ-treatment, Tg(*flt4:mCitrine*) larvae were transferred to a solution containing 10mM pentylenetetrazol (PTZ) (Sigma-Alrich, UK) dissolved in E3 media or the same volume of E3 media for 1 hr (ZT1-2) as previously described (Reichert et al, 2019). PTZ-treated fish were assessed for behavioural hyperactivity and tachycardia using a Nikon SMZ1500 bright-field microscope, subsequently washed out of solution 3x in E3 media, anaesthetised in MS222 (42 mg/L) and intraventricularly injected (see *injection regime*) and imaged for several hours (see *live imaging*). For melatonin-treatment, Tg(*flt4:mCitrine*) larvae were intraventricularly injected at ZT1 and immediately transferred into solution containing 1µM melatonin (Sigma-Alrich, UK) dissolved in 0.2% DMSO or 0.2% DMSO control for subsequent imaging. For caffeine-treatment, Tg(*flt4:mCitrine*) larvae were intraventricularly injected at ZT19 and immediately transferred into solution containing 250µM caffeine (Sigma-Alrich, UK) dissolved in E3 media or E3 media alone for subsequent imaging. For inclusion and filopodia analysis, the treatment group was placed in melatonin solution (1 µM) or caffeine solution (250µM) and control group in 0.2% DMSO or H20 respectively, for 4 hrs (ZT1-5 for melatonin and ZT-19-23 for caffeine) before they were culled, fixed in 4%PFA and imaged (see *muLEC cell and subcellular morphology)*.

#### Sleep deprivation assay

Sleep deprivation by gentle handing was performed as previously described (Suppermpool et al., 2024). Prior to lights-off (ZT14), *Tg*(*flt4:mCitrine*) larvae were transferred into two separate identical 24-well clear plates (ThermoFisher, UK). Immediately after lights off (ZT14-18) both plates were placed in a dark room under dim-red light (Blackburn Local Bike Rear Light 15 Lumen, UK). Larvae in one plate were left undisturbed while larvae in the other plate were manually sleep deprived by repeated gentle stimulation using a No. 1-2 paintbrush (Daler Rowney Graduate Brush, UK) to prevent larvae from being immobile for longer than 1 minute. Placing the paintbrush into the water of the well was usually stimulatory enough to elicit movement, but if larvae remained immobile, they were gently touched until movement was initiated. The 4-hr sleep deprivation was performed by two experimenters in 2-hr shifts.

#### Injection regime

Intraventricular injections were conducted as previously described (Shibata-Germanos et al, 2020) using a PM1000 Cell Microinjector (MicroData, Instrument, Inc) with glass capillary needles (Harvard Apparatus, # GC100F-15) prepared with a Micropipette Puller (Shutter Instruments). Larvae were anaesthetised in E3 media with MS222 (42 mg/L) and mounted dorsally in 2% low-melt agarose (Sigma Aldrich, UK) dissolved in ddH_2_0 on a 60mm plastic petri dish lid which was scratched with forceps prior to mounting to reinforce agarose adherence. The following substrates were injected into the rhombenchephalic ventricle with a total bolus volume 1nl at 20 psi pressure at the specified concentration: Dextran, Fluorescein 2MDa, 1mg/ml (ThermoFisher, UK); pHrodo Red Avidin, 2mg/ml (ThermoFisher, UK); pHrodo Red Dextran 10kDa, 0.5mg/ml (ThermoFisher, UK); pHrodo Red LDL, 1mg/ml (ThermoFisher, UK). pHrodo-conjugated dyes were pre-mixed with Dextran, Fluorescein 2MDa immediately prior to co-injection so that in each 1nl bolus, 0.5nl constituted each respective dye.

#### Live imaging

All live imaging data were acquired using a Zeiss LSM 980 confocal microscope with a 20x/1.0 NA water immersion objective. Z-stacks were obtained with sequential acquisition settings of 1024 × 1024 pixels. Z-depth intervals ranged from 1-1.5μm which were standardised within each assay. Dextran, Fluorescein 2MDa was excited at 488nm and pHrodo Red Avidin, pHrodo Red Dextran and pHrodo Red LDL were excited at 546nm. Multi-position time lapses were set up and larvae were sequentially imaged every 60 minutes post-injection over the course of 6-7 hours, unless otherwise stated.

#### Image Processing

Confocal stacks were processed and analysed in Imaris. Briefly, raw z-stacks were segmented and thresholded to provide representative surfaces of a given object in 3D space. Segmentation and thresholding limits were standardised across all fish in a given assay. Standardised regions of interest (ROIs) were constructed for muLECs along the midline of both tectal rings as representative to reduce variability of data from lateral muLECs at lower z-depths. For assessment of the initial injection bolus, 3D surfaces of Dextran, Fluorescein 2MDa signal were constructed using standardised ROIs spanning the most anterior point of the telencephalic ventricle to the posterior point of the diencephalic ventricle. All z-stacks are represented as dorsal maximum projections unless otherwise stated.

### Quantitative immunohistochemistry and imaging

Larvae were culled by anaesthetic overdose (MS222, approximately 420 mg/L) and fixed overnight (12-16 hours) in 4% PFA supplemented with 4% sucrose at 4°C. The following day, larvae were washed three times in 1x PBS and dissected. To remove batch effects, day and night larvae were given distinctive oblique cuts on the tail for later identification and transferred to the same vial for staining. Dissected larvae were permeablised with Proteinase K (30 μg/mL) for 90 seconds, post-fixed for 20 minutes in 4% PFA at room temperature and subsequently washed three times in PBST. Samples were blocked with blocking buffer (10% normal goat serum; 1% DMSO; 0.5% Triton-X100 in PBS) for 1 hr at room temperature. Samples were then incubated with a custom anti-Mrc1a antibody (1:200 dilution, rabbit polyclonal, Sino Biological, UK) in blocking buffer overnight at room temperature. Primary antibody was removed, and samples were rinsed 3x in PBST and washed 4×30mins in PBST. Samples were placed in fresh block buffer with 1:500 anti-rabbit Alexa Fluor 568 (ThermoFisher, UK). Excess secondary antibodies were washed 4-6 times for 30 minutes and mounted dorsally in 2% LM agarose for imaging.

All imaging data were acquired using a Zeiss LSM 980 confocal microscope with a 20x/1.0NA water-immersion objective. Z-stacks were obtained at 1.5μm depth intervals with sequential acquisition settings of 1024 × 1024 pixels. Endogenous mCitrine and anti-rabbit Alexa Fluor 568 were excited at 488nm and 561nm, respectively. Image analysis was conducted in Imaris. Raw z-stacks were segmented and standardised thresholds were applied to provide representative 3D surfaces of muLEC and anti-Mrc1a immunofluorescence.

### Generation of *mrc1a* F_0_ knockouts

#### crRNA Selection

F_0_ knockout larvae were generated as described previously (Kroll et al., 2021; Kroll et al., 2025). All crRNAs used in F_0_ knockout experiments were designed using CHOPCHOP (chopchop.cbu.uib.no) (Labun et al., 2019). Putative targets were checked for SNPs using SNPfisher (Butler et al., 2015). Sequences of the targets (and genomic positions) used to generate F_0_ knockout are as follows with the protospacer adjacent motif (PAM) underlined (5’ 3’): mrc1a_guide_1, TGAGGGTCAACGCTTTCGAT<u>GGG</u> (chr7:58939687); mrc1a_guide_2, ACTACGGAGGGAAGATCAGA<u>TGG</u> (chr7:58938982); mrc1a_guide_3, TGGGACAGTGATCCAGTGAC<u>TGG</u> (chr7:58936563). Scrambled, nonsense crRNAs were prepared identically and had the following sequences (5’ 3’): scrambled_guide_1, CCCGACTAACATGGCGAGAC; scrambled_guide_2, GTATCCACCACGGGAGACAC; scrambled_guide_3, GTATATGGTTTGGGGCCACT; scrambled_guide_4, CGCAAGTTAAGGCCGACCCA. Genomic positions are for reference genome GRCz11 (danRer11).

#### Cas9/gRNA preparation

Cas9/gRNA were prepared as previously described (Kroll et al., 2021; Kroll et al., 2025). CRISPR-Cas9 RNPs were composed of: crRNA (Alt-R® CRISPR-Cas9 crRNA) and tracrRNA (Alt-R® CRISPR-Cas9 tracrRNA), Cas9 (Alt-R® S.p. Cas9 Nuclease V3) (Integrated DNA Technologies, UK). crRNA and tracrRNA were resuspended in Duplex buffer (IDT, UK) to form 200 μM stocks. Stocks of crRNA and tracrRNA were stored at −70°C before use and Cas9 was stored at –20°C before use. Each crRNA was annealed separately with the tracrRNA by mixing 1 μL crRNA 200 μM; 1 μL tracrRNA 200 μM; 1.28 μL Duplex buffer. The mix was heated to 95°C for 5 min, then cooled on ice, to obtain a 61 μM gRNA solution. The gRNA solutions were then mixed in equal volumes with Cas9 (1 μL gRNA 61 μM; 1 μL Cas9 61 μM directly from the IDT vial), incubated at 37°C for 5 min then cooled on ice, generating three 30.5 μM RNP solutions. The three RNP solutions were pooled; the final concentration of each RNP in the pool was thus 10.2 μM and the total RNP concentration 30.5 μM. The RNPs were usually kept overnight in a 4°C fridge on ice before injections the following day. Some experiments used RNPs stored at –70°C for a few weeks.

#### Injections

Approximately 1nL of the three-RNP pool (∼10.2fmol per RNP) was injected into the yolk at the single-cell stage before cell inflation. Injected embryos were kept at room temperature for approximately 45 minutes post-injection before transfer to a 28.5°C incubator, as delaying the first cell divisions may increase mutagenesis and reduce the diversity of alleles (Terzioglu et al., 2020).

#### Preparation of samples for Illumina MiSeq

Target loci were deep sequenced by Illumina MiSeq as previously described (Kroll et al., 2021; Kroll et al., 2025). 8dpf *mrc1a* KO and scrambled control larvae were culled by MS222 overdose and genomic DNA was extracted by HotSHOT as previously described (Meeker et al., 2007). Individual larvae were transferred to a 96-well PCR plate where excess liquid was removed from each well before adding 50 μl of 1x base solution (25 mM KOH, 0.2 mM EDTA in water). Plates were sealed and incubated at 95°C for 30 min then cooled to room temperature before the addition of 50 μL of 1x neutralisation solution (40 mM Tris-HCl in water). Genomic DNA was then stored at –20°C. PCR primers were designed for each target locus using Primer-BLAST (NCBI) to amplify a window of 150–200 bp with at least 30 bp between each primer binding site and the predicted double-strand break site. Forward and reverse PCR primers were ordered with overhang sequences at the 5′-end and 3’ end of each primer to allow indexing (Forward 5’ overhang: TCGTCGGGCAGCGTCAGATGTGTATAAGAGACAG; Reverse 3’ overhang: GTCTCGTGGGCTCGGAGATGTGTATAAGAGACAG).

Each PCR well contained: 7.98 μL PCR mix (2 mM MgCl_2_, 14 mM pH 8.4 Tris-HCl, 68 mM KCl, 0.14% gelatine in water, autoclaved for 20 min, cooled to room temperature, chilled on ice, then added 1.8% 100 mg/mL BSA and 0.14% 100 mM d[A, C, G, T]TP), 3 μL 5× Phusion HF buffer (New England Biolabs, UK), 2.7 μL dH2O, 0.3 μL forward primer (100 μM), 0.3 μL reverse primer (100 μM), 0.12 μL Phusion High-Fidelity DNA Polymerase (New England Biolabs, UK), 1.0 μL genomic DNA; for a total of 15.4 μL. The PCR plate was sealed and placed into a thermocycler. The PCR program was: 95°C – 5 min, then 40 cycles of: 95°C – 30 sec, 60°C – 30 sec, 72°C – 30 sec, then 72°C – 10 minutes, then cooled to 10°C until collection. The PCR product’s concentration was quantified with Qubit (dsDNA High Sensitivity or Broad Range Assay) and its length was verified on a 1.5% agarose gel with GelRed (Biotium, USA). Excess primers and dNTPs were removed by ExoSAP-IT (ThermoFisher, UK) following the manufacturer’s instructions. The samples were then sent for Illumina MiSeq, which used MiSeq Reagent Nano Kit v2 (300 Cycles) and analysed as in Kroll et al, 2025.

#### Morphological assessments of mrc1a F_0_ knockouts

*mrc1a* KOs and scrambled controls were generated in the following lines: *Tg*(*flt4:mCitrine*), *Tg*(*flt4:mCitrine);Tg(kdrl:mCherry*), and *Tg*(*flt4:mCitrine); Tg(mpeg:mCherry*). At 5dpf, larvae were anaesthetised (MS222, 42 mg/L), mounted dorsally in 2% LM agarose on a small petri dish lid. Imaging data were acquired using a Zeiss LSM 980 confocal microscope with a 20x/1.0NA water immersion objective. Z-stacks were obtained with sequential acquisition settings of 1024 × 1024 pixels with z interval depths of 1μm. mCitrine and mCherry were excited at 488nm and 594nm, respectively.

Confocal stacks were processed and analysed in Imaris. For muLEC morphology assessment, standardised ROIs were constructed bilaterally over the optic tectum and standardised threshold and segmentation were applied to construct 3D surfaces using the Imaris Surface function. For assessment of the vascular network, as shown previously (Spangenberg et al, 2023), standardised ROI were constructed from the dorsal midline junction to the most anterior point of the prosencephalic artery to depth of 120μm from the skins surface as visible by autofluorescent signal. The Imaris Filament function with loops based on local intensity contrast without the detection of spines was performed to reconstruct the vasculature within the ROI as a connected network in 3D space. For assessment of *mpeg*^+^ cells, the Imaris Surfaces and Filament (with tree autopath algorithm) function was used to reconstruct mCherry positive signal within a standardised ROI covering the head region and eyes at a depth of 120μm from the autofluorescent skin into 3D space.

### muLEC ablation and seizure recovery

#### muLEC ablation

*Tg(flt4:mCitrine); Tg(kdrl:mCherry)* larvae were raised to 5dpf, anaesthetised in 42 mg/L MS222 solution and dorsally mounted in 2% LM agarose. mCitrine positive cells within the bilateral muLEC tectal loops and along the midline were ablated using a Zeiss LSM 980 confocal microscope with two-photon laser at 800nm (Mai Tai HP laser 690-1040nm tuneable, Spectra Physics). Control ablations were performed by directing an identical laser pulse just adjacent to mCitrine positive muLECs in a fluorescent negative space. Larvae were imaged before and after muLEC-ablation and mock-ablation to confirm successful ablations and intact surrounding blood vasculature. Larvae were re-imaged 3 days post ablation to assess for regeneration. Regenerated muLECs were quantified in Imaris using the Imaris Surface function.

#### PTZ-induced seizures and locomotor tracking

Following ablations, larvae were unmounted from agarose and transferred to a solution 10mM PTZ dissolved in E3 media or the same volume of E3 media alone for 1 hr at approximately ZT7. PTZ-treated fish were assessed for behavioural hyperactivity and tachycardia using a Nikon SMZ1500 bright-field microscope, subsequently washed out of solution 3x in E3 media and transferred into individual wells of a 96-well plate (ThermoFisher, UK) containing approximately 600μL of fish water for behavioural tracking.

Locomotor activity of larvae was monitored using an automated video tracking system (Zebrabox, Viewpoint LifeSciences) in a temperature-regulated room (26.5°C) and illuminated with constant white light. Larval movement was recorded using the Videotrack ‘quantization’ mode with the following detection parameters: detection threshold, 15; burst, 100; freeze, 3; bin size, 60s. Tracking was performed at 25 frames per second for approximately 48 hours. Behavioural data was analysed with custom Matlab (R2024a, the Mathworks) scripts (available at https://github.com/JRihel/muLEC-Circadian-Imaging).

### Data Analysis and Statistical tests

Imaging processing, analysis and presentation was performed in Imaris 9.82 - 10.0.0 (Imaris, UK) and Fiji-Image J version 2.14.0i. Graph construction and statistical analyses were performed in Prism 9.2.0 (GraphPad software, USA). All data were assessed for normality by a Shapiro-Wilk test. Parametric data were assessed by One-way ANOVA (two-tailed), Mixed-Effects Model and log rank tests, and non-parametric data were assessed by Mann-Whitney U tests (two tailed) as specified. All tests were conducted with an alpha of 0.05. Figures and schematics were constructed in Inkscape version 1.4.

---

**Figure S1:**
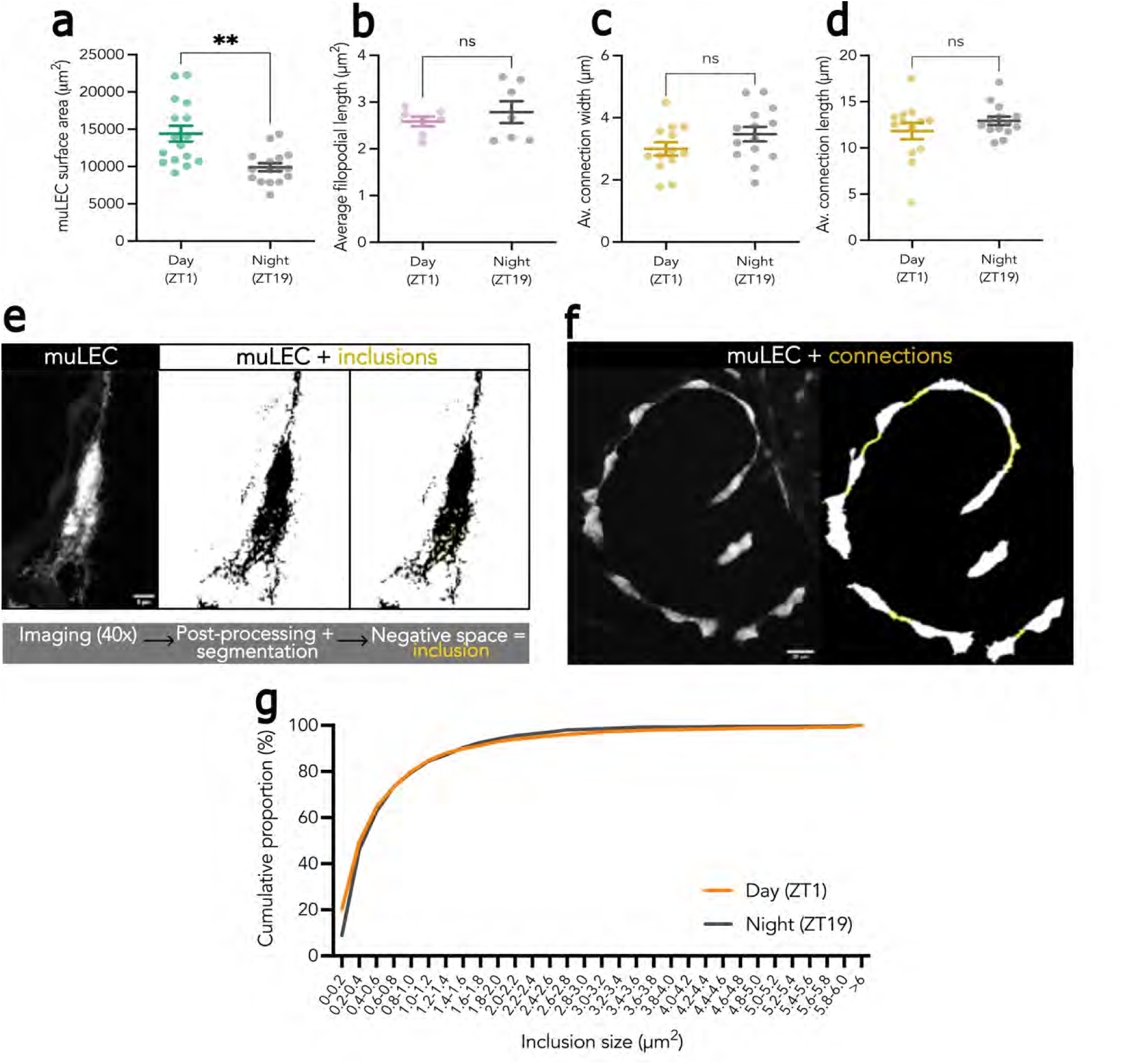
Additional morphological characteristics of muLECs over the day night cycle and analysis methodology. **a)** Surface area of the muLEC loop day (n=16) vs night (n=16). **b**) Length of muLEC filopodial protrusions in the day (n=7) and night (n=7). **c,d**) The average width (c) and length (d) of connections between muLECs between the day (n=13) and night (n=14). **e**) Confocal imaging of a single muLEC in a *Tg(flt4:mCitrine)* larva (grayscale) was processed and segmented in 2D to identify fluorescent-negative spaces as intracellular inclusions. **f)** Morphometric analysis of the muLEC tectal loop (grayscale) reveals connections (yellow) between muLECs. **g**) Distribution of inclusion size as a cumulative percentage frequency of total recorded inclusions in all cells in all larvae during the day versus night. **a-d**, data are mean ± SEM; each dot represents one larva. Statistical differences were determined by one-way ANOVA (two-tailed, **a-d**). ^ns^ p>0.05; *** p<0.001. Scale bar: 5µm (e); 20µm (f).

**Figure S2:**
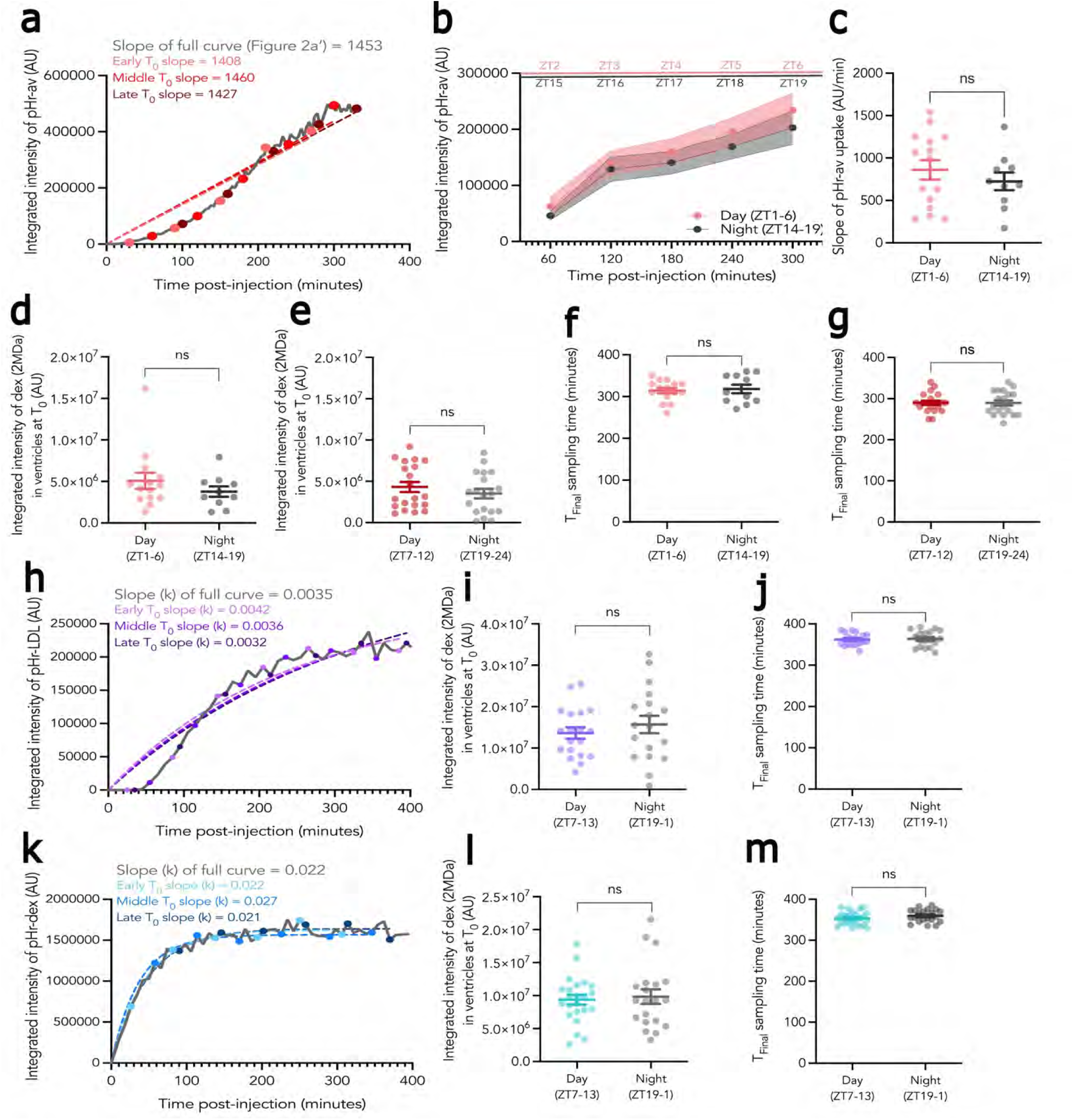
muLEC-uptake curve modelling and experimental controls. **a)** The rate of muLEC-uptake of pHrodo-avidin is well captured by a simple linear regression when data are sampled at low frequency (every 60 min), with the slope from the linear fit similar to 4 minute sampling (grey continuous line; data from Figure 2b) irrespective of early, middle or late starting points (T0= 30, 60, 90 mins post-injection, respectively). **b**) The average integrated intensity ± SEM of pHrodo-avidin uptake by muLECs is similar between the early day (ZT1-6, n=15) and early night (ZT14-19, n=11). **c**) Slope of muLEC-uptake of pHrodo-avidin from a simple linear regression. **d,e)** Average integrated intensity at T0 of 2MDa dextran across both telencephalic and diencephalic ventricles is the same across groups presented in b,c (d) or Figure 2e,f (e). **f,g**) Time-point Tfinal is the same between groups presented in b,c (f) or Figure 2e,f (g). **h**) Representative larva sampled every 10 minutes post-injection of pHrodo-LDL (grey line). One-phase association regression captures slopes when simulated at low frequency. **i,j**) There is no difference in injection bolus (**i**) or time-sampling (**j**) from the data presented in Figure 2h,i. **k**) Representative larva sampled every 8 minutes post-injection of pHrodo-dextran (grey line). One-phase association regression captures slopes when simulated at low frequency. **l,m**) There is no difference in injection bolus (l) or time-sampling (m) from the data presented in Figure 2k,l. Data are mean ± SEM and each dot represents one larva. Statistical differences were determined by one-way ANOVA (two-tailed, **b-e**, **g-k**). ^ns^ p>0.05.

**Figure S3:**
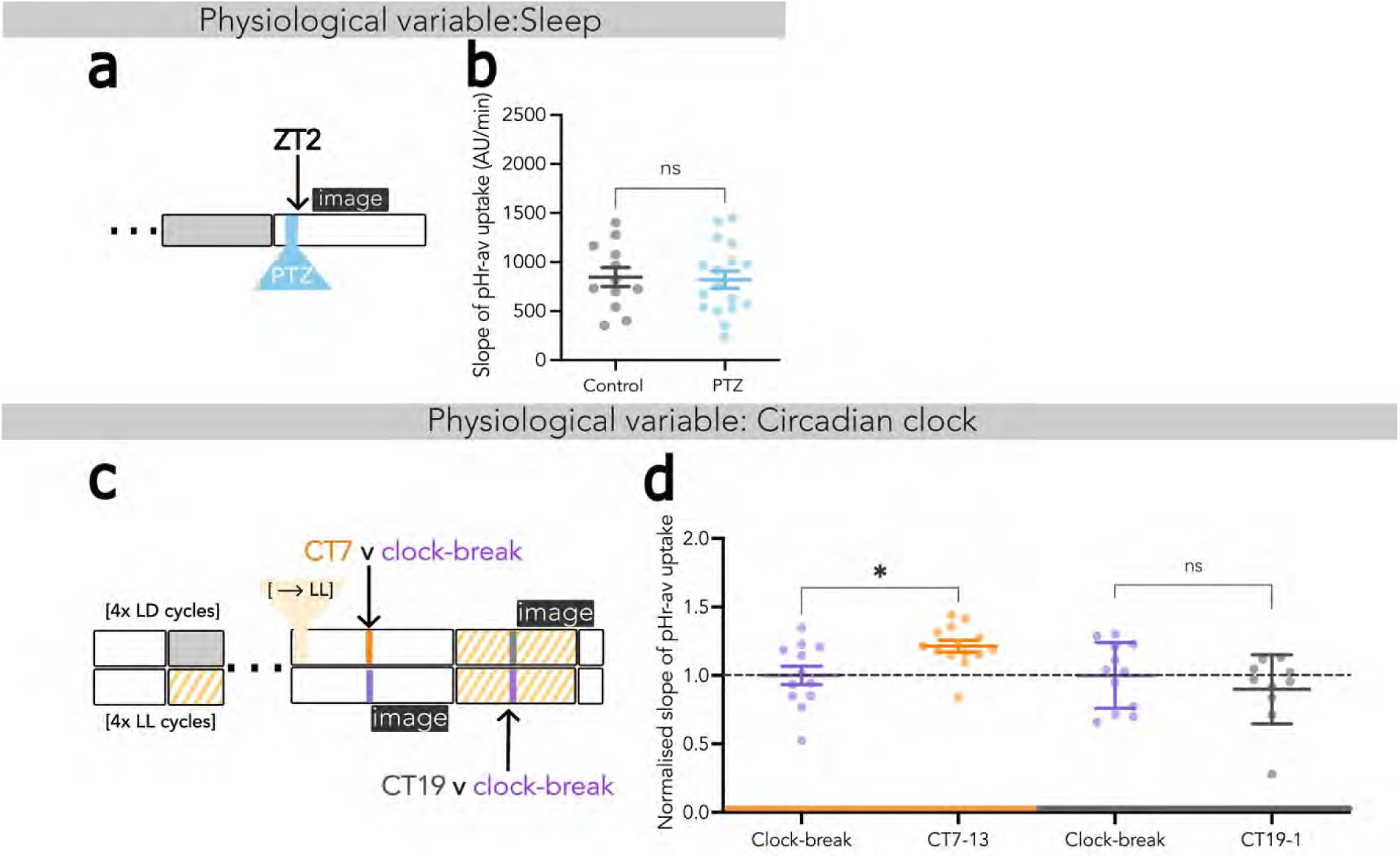
Diurnal variation of muLEC-uptake of pHrodo-avidin is regulated by the clock and not brain state. a) Experimental design to test the effect of drug-induced rebound sleep. **b**) The slope of pHrodo-avidin uptake was not significantly different between PTZ treated (n=17) and controls (n=12). **c**) *Tg(flt4:mCitrine)* embryos were either (top) entrained on a normal light-dark cycle to 4dpf before switching to free-running conditions (as in Figure 2b) or (bottom) raised under constant light conditions to suppress circadian rhythm development, referred to as ‘clock-break’. **d**) Normalised slope of muLEC-uptake of pHrodo-avidin from a simple linear regression revealed that entrained larvae show higher levels of muLEC-uptake relative to clock-broken fish (purple) when sampled in the middle of their subjective day (CT7, yellow) and not in the middle of their subjective night (CT19, grey). As per x-axis: Clock break, n=12; CT7, n=14; Clock break, n=12; CT19, n=10. Data are mean ± SEM; each dot represents one larva. Statistical differences were determined by one-way ANOVA (two-tailed). ^ns^ p>0.05; * p<0.05.

**Figure S4:**
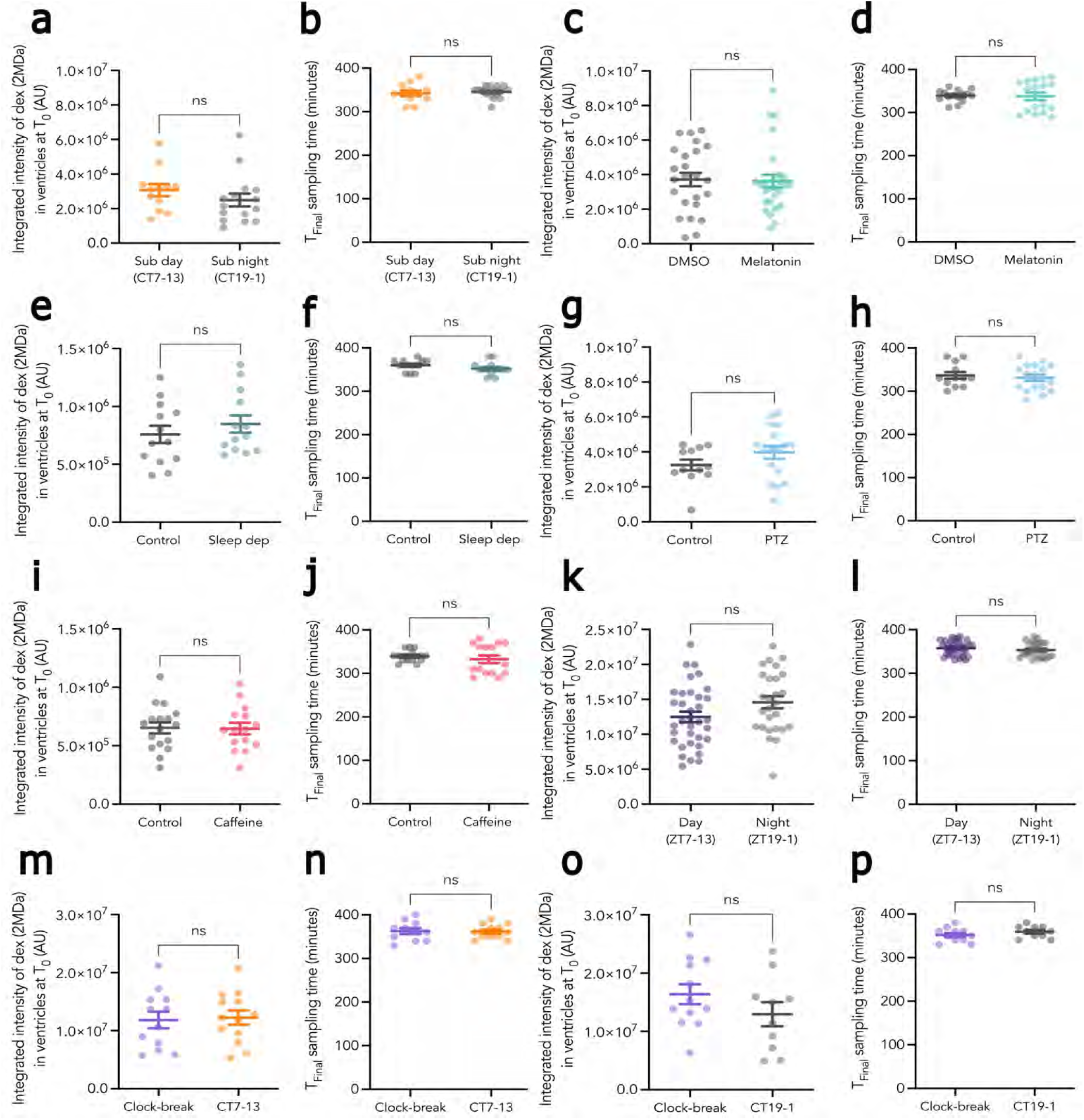
Injection dose and final sampling times for Figure 3 and Figure S3. Average integrated intensity of 2MDA dextran signal in both the telencephalic and diencephalic ventricles at T0 is the same between groups corresponding to experiments presented in: **a**, Figure 3c; **c**, Figure 3e; **e**, Figure 3g; **g**, Figure S3b; **i**, Figure 3i; **k**, Figure 3m,n; **m,o**, Figure S3d. Final sampled time-points post-injection demonstrating no sampling bias between groups corresponding to experiments presented in: **b**, Figure 3c; **d**, Figure 3e; **f**, Figure 3g; **h**, Figure S3b; **j**, Figure 3i; **l**, Figure 3m,n; **n,p**, Figure S3d. All data are mean ± SEM; each dot represents one larva. Statistical differences were determined by Mann-Whitney U tests (two-tailed, a,b,d,f,j); one-way ANOVA (two-tailed, c,e,g,h,l,k-p). ^ns^ p>0.05.

**Figure S5:**
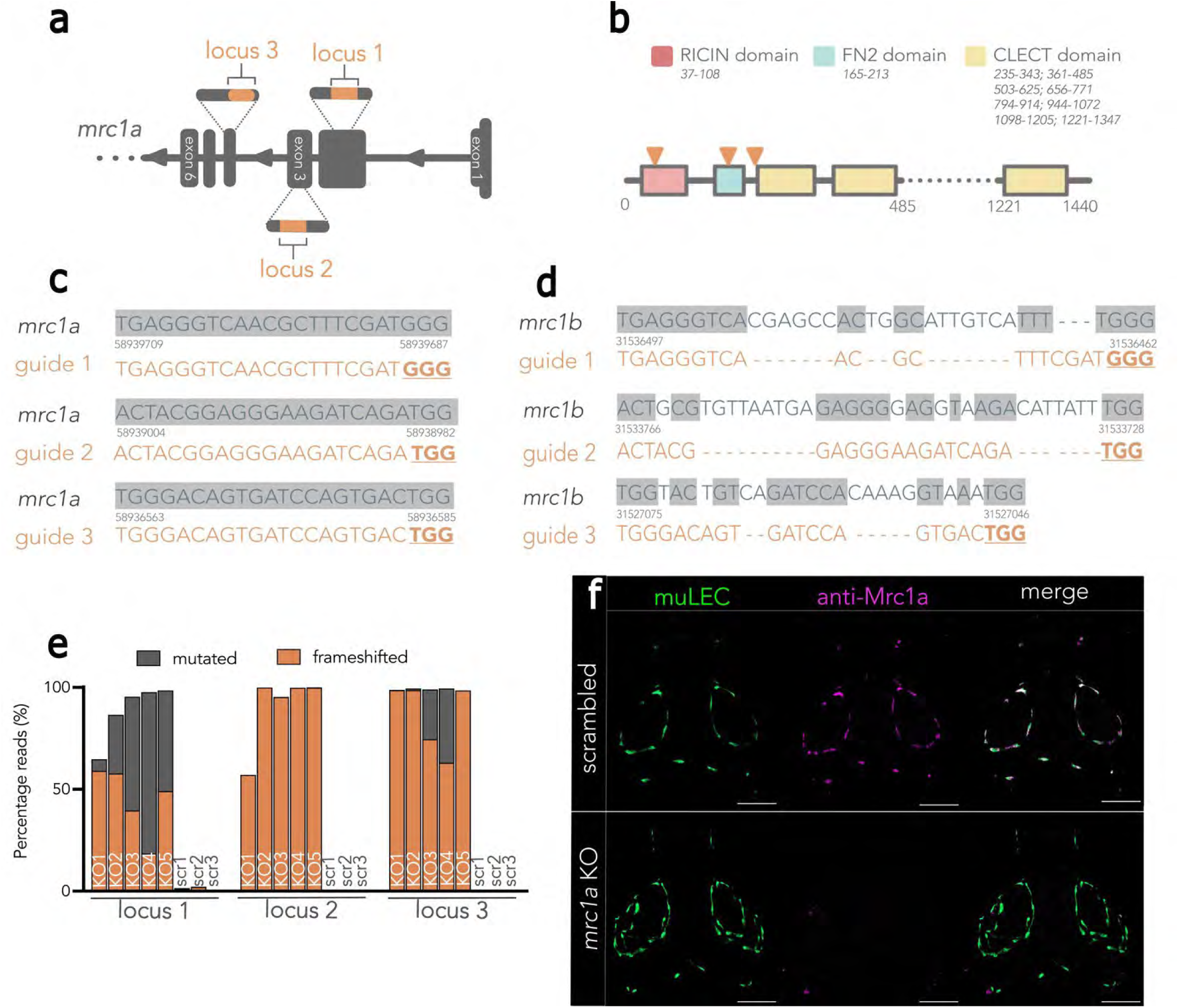
Deep sequencing of *mrc1a* F0 knockouts and validation of the anti-Mrc1a antibody. a) Schematic of the *mrc1a* gene in the 5’-3’ genomic direction. Exons are displayed in dark grey; short exons are protein coding; tall exons are 5’ or 3’ UTRs. Orange lines show the 3 target loci in exons 2, 3 and 4 for CRISPR-mediated mutagenesis of *mrc1a*. Exons and introns are not to scale. **b**) Schematic of the Mrc1a protein with key conserved protein domains and their respective amino acid positions. Orange arrows indicate the predicted locations of truncations generated from the 3 target loci. **c**, **d**) Genomic sequences of *mrc1a* (c) and *mrc1b* (d) showing that the selected gRNAs targeted *mrc1a* and not *mrc1b*. Genomic bases complimentary to the guide have a grey background. The protospacer adjacent motif (PAM) where Cas9 binds is underlined in orange. **e**) Stacked plot of the percentage of total reads mutated (grey) and with a frameshift mutation (orange) as computed by ampliCan analysis following MiSeq deep sequencing of each targeted locus of *mrc1a* (white label, knockout, n=5) or in scrambled controls (grey label, scr, n=3). Each number represents the same larva sequenced at multiple loci. **f**) Dorsal maximum projection of anti-Mrc1a immunohistochemistry (magenta) and endogenous fluorescence of muLECs (green) in 5dpf *Tg(flt4:mCitrine)* larvae. Mrc1a immunofluorescence is present in scrambled controls (top) but lost in *mrc1a* KO larvae (bottom). Abbreviations: RICIN, ricin B, lectin domain; FN2, fibronectin type II; CLECT, C-type Lectin; Scale bar, 100µm.

**Figure S6:**
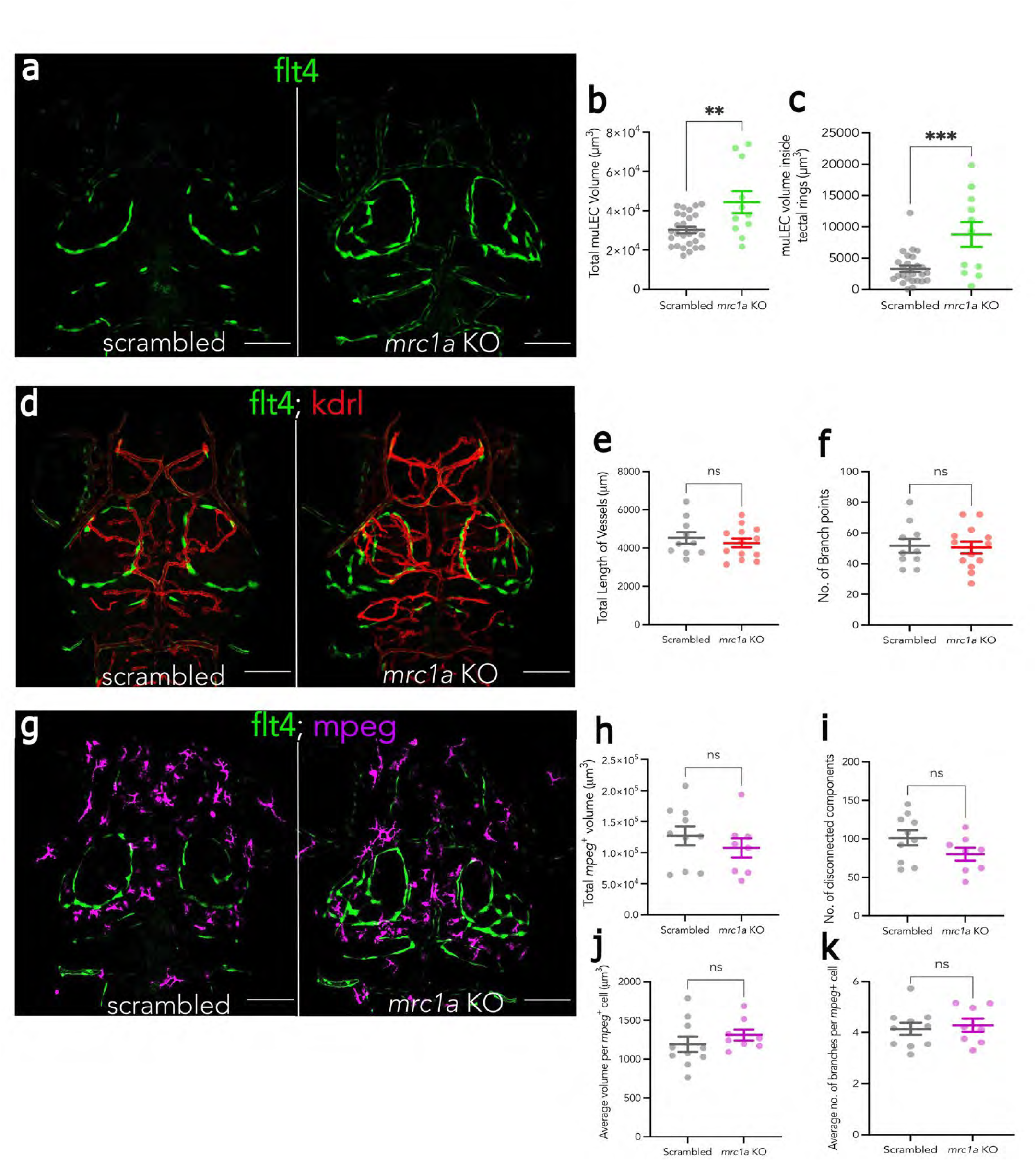
*mrc1a* knockout affects muLEC morphology but not the surrounding vasculature or *mpeg+* macrophages. *mrc1a* KO or control larvae were generated in 3 transgenic lines that label **a**) muLECs (Tg(*flt4:mCitrine*)) in green; **d**) muLECs and surrounding vasculature in red (*Tg*(*flt4:mCitrine*)*;Tg*(*kdrl:mCherry*)); and **g**) muLECs and *mpeg*+ cells in magenta (*Tg*(*flt4:mCitrine*)*;Tg(mpeg:mCherry*)). Representative scrambled-control larva (left) and *mrc1a* F0 KO larva (right) are shown. **b**,**c**) muLEC morphology is quantified as volume of mCitrine+ mass in the bilateral loops over the optic tectum (**b**) or populating the inner tectal rings (**c**) reveals altered muLEC morphology in *mrc1a* F0 KOs (green, n=11) compared to scrambled-injected controls (grey, n=27). **e**,**f**) Vascular morphology is quantified in *mrc1a* KOs (red, n=13) or controls (grey, n=10) by (**e**) the cumulative length of the *kdrl*+ vessels in diencephalic and telencephalic regions 120μm from the skin surface and (**f**) the total number of branching points. **h-k**) There were no differences between *mrc1a* KOs (magenta, n=8) and controls (grey, n=10) in total mpeg+ cell volume in the head cavity (**h**), or the number (**i**), volume (**j**) or branches (**k**) of individual *mpeg*+ cells. Data are mean ± SEM; each dot represents one larva. Statistical differences were determined by one-way ANOVA (two-tailed). ^ns^ p>0.05; ** p<0.01; *** p<0.001. Scale bar:100µm.

**Figure S7:**
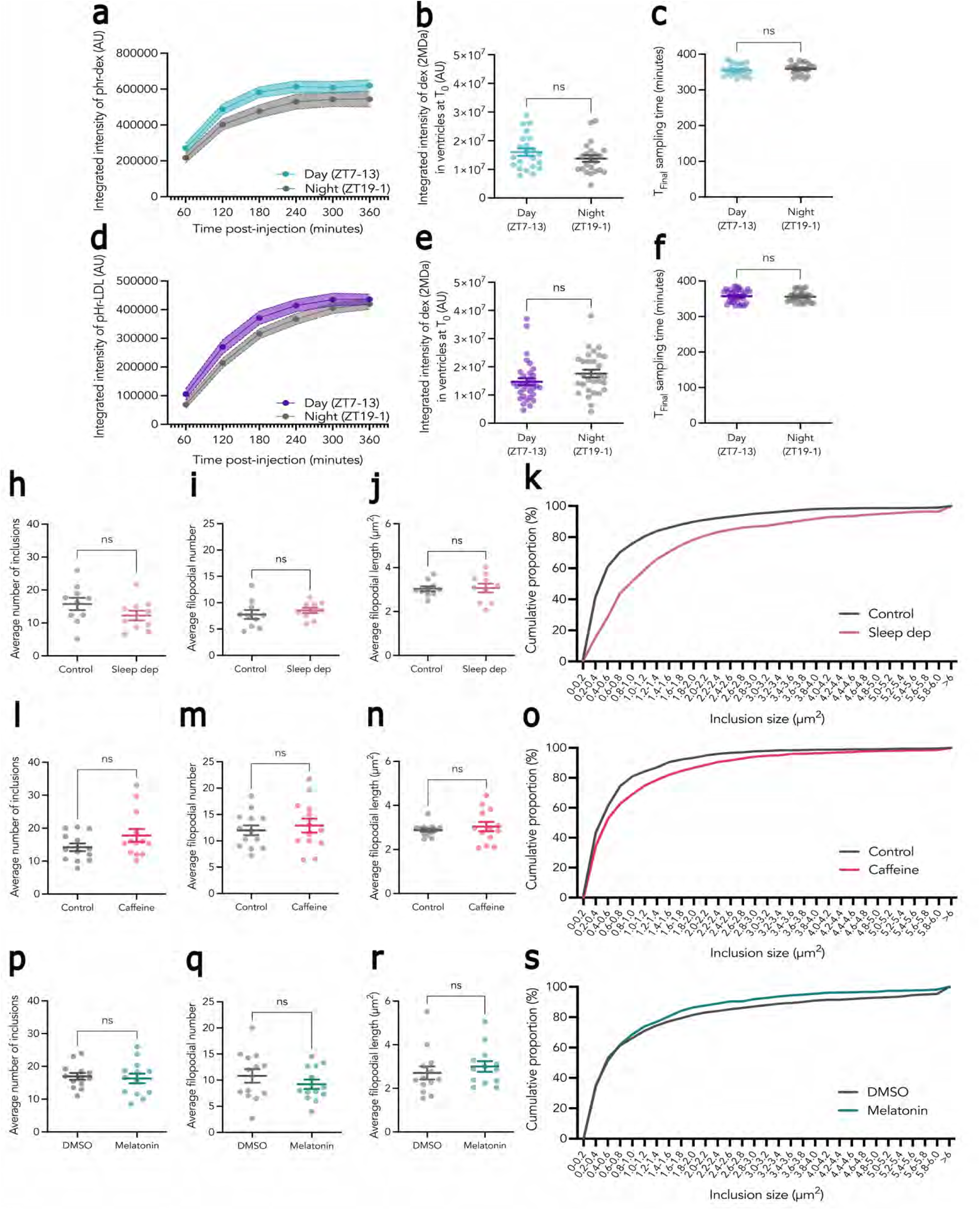
supplemental data relating to Figure 5. **a**, **d)** The average integrated intensity ± SEM of pHrodo-dex (a) or pHrodo-LDL (d) accumulated in muLECs in *Tg(flt4:Gal4);Tg(UAS:GFP2AdnCLK)* larvae day (ZT7) vs night (ZT19). Data corresponding to Figure 5b (a) and Figure 5c (d). **b,e**) Average integrated intensity of 2MDa dextran signal in both telencephalic and diencephalic ventricles at T0 is not different between groups, corresponding to experiments presented in Figure 5b (b); Figure 5c (e). **c,f**) Time Tfinal is similar between groups corresponding to experiments presented in: Figure 5b (c); Figure 5c (f).**h,l,p**) Average number of intracellular muLEC inclusions per larvae following: h, sleep deprivation (n=10) or undisturbed sleep (n=10); l, Caffeine-treatment (n=13) or fish water control (n=13); p, Melatonin-treatment (n=13) or DMSO controls (n=13). **i,j,m,n,q,r**) Average number and length of filopodia per cell per larva following: i,j, sleep-deprivation; m,n, Caffeine-treatment; q,r, Melatonin-treatment. **k,o,s**) Distribution of inclusion size as a cumulative percentage frequency of all recorded inclusions in all cells in all larvae following: k, sleep-deprivation; o, Caffeine-treatment; s, Melatonin-treatment. Data are mean ± SEM; each dot represents one larva (b,c,e,f,h-j,l-n,p-r). Statistical differences were determined by Mann-Whitney U tests (two-tailed, b,c,p,r), or one-way ANOVA (two-tailed, e,f,h,i,j,m,n,q). ^ns^ p>0.05.

**Figure S8:**
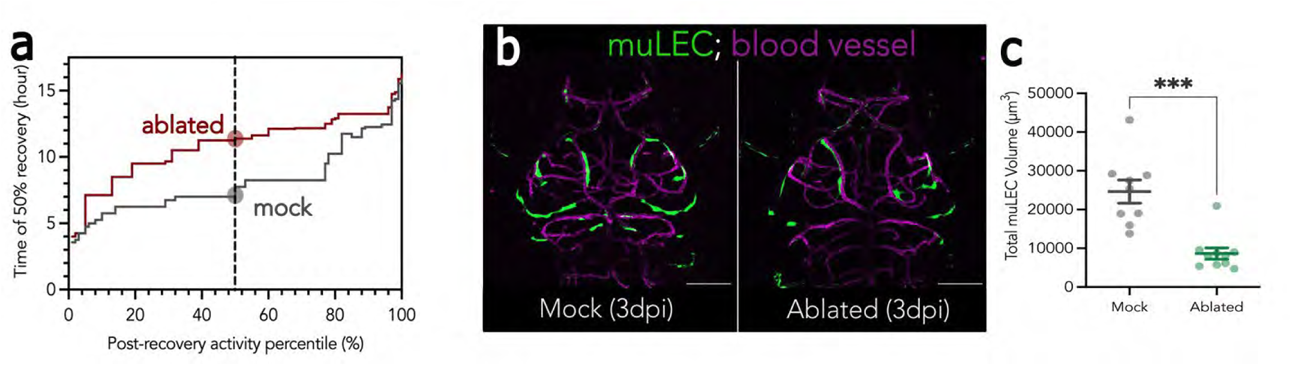
Behavioural recovery post treatment and muLEC regeneration post-ablation. **a**) The delay in the recovery time (i.e., when 50% of larvae have recovered) in muLEC-ablated fish is observed across a wide range of recovery cutoff values (Activity Percentiles). The values derived from the curves in Figure 6c are noted by the vertical dashed line. **b**) Representative dorsal maximum projections of a mock-ablated (left) and muLEC-ablated (right) larva 3 days post intervention (dpi). muLECs (green) and blood vessels (magenta) in a (*Tg(flt4:mCitrine;Tg(kdrl:mCherry larva)*. **c**) Regenerative capacity of muLECs post-ablation as quantified by the volume of mCitrine+ cells in bi-lateral tectal loops structures 3 days post-muLEC (n=10) or mock ablation (n=9). Data are mean ± SEM; each dot represents one larva. Statistical differences were determined by Mann-Whitney U tests (two-tailed, c). ^ns^ p>0.05; *** p<0.001.

**Supplemental Video 1:**
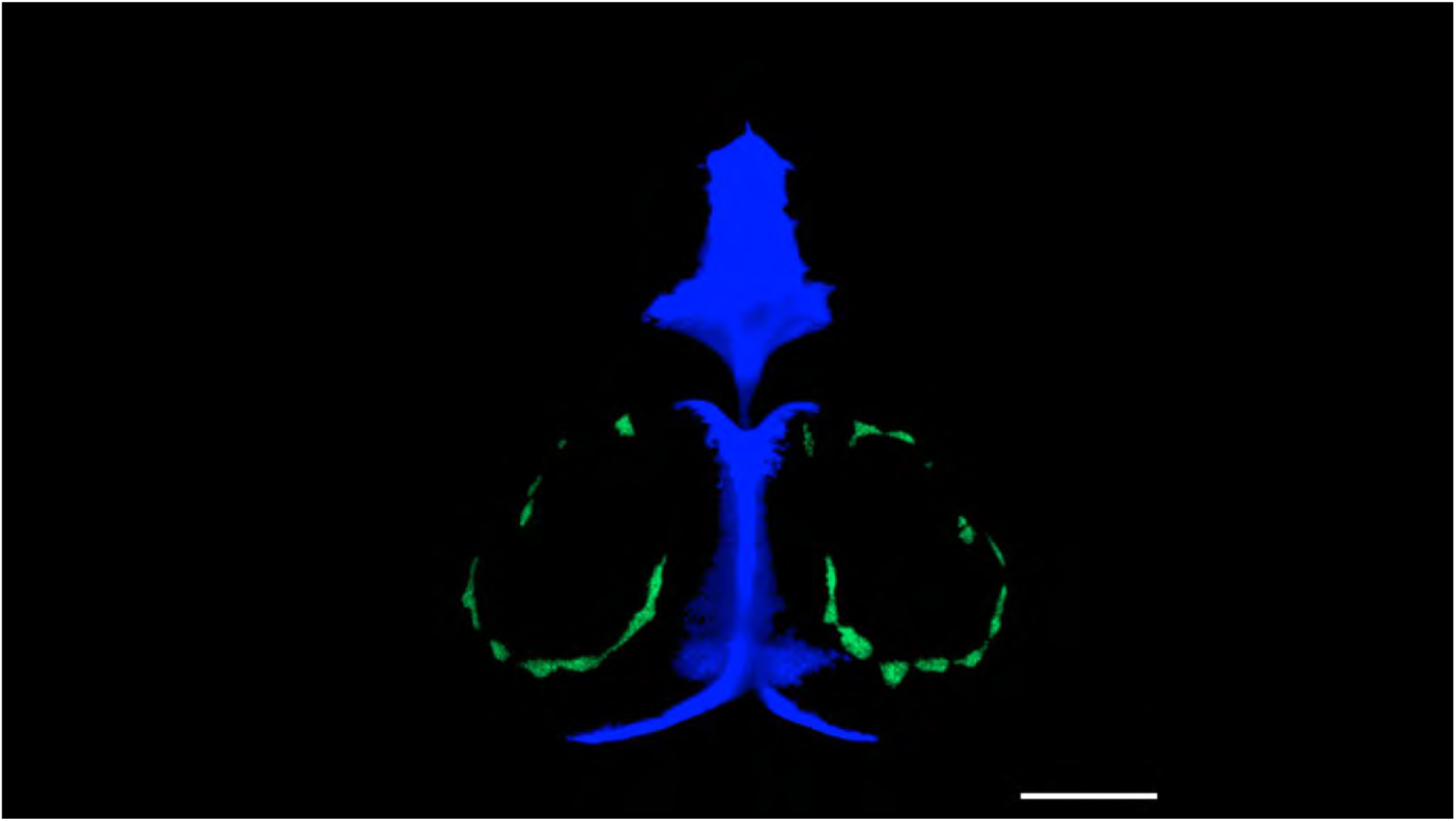
Timelapse of a 5dpf *Tg(flt4:mCitrine)* larva injected with 2MDa Alexa-fluor, 488 dextran (Dex; blue) and pHrodo Red avidin (pHr-av; red) and imaged at high-time resolution as depicted in Figure 2a, S2a. Signal was extracted from muLEC and ventricle masks removing any brain derived signal. First frame captured 5 minutes post-injection and every subsequent frame +4 minutes for a total imaging window of 330 minutes. Scale bar, 100µm.

